# Lab-on-a-3D-Printer for Democratized Reconfigurable Digital Microfluidics

**DOI:** 10.64898/2026.01.01.697313

**Authors:** Shun Ye, Bella Rose Schremmer, Zixin Guan, Artem Goncharov, Gyeo-Re Han, Vivek Rajasenan, Yichen Zou, Xiang Li, Charlotte Rose McDonough, Aydogan Ozcan, Dino Di Carlo

## Abstract

Automation in the life sciences remains dominated by expensive, centralized robotic systems, leaving most laboratories unable to access precision robotic liquid handling. Here we present the Lab-on-a-3D-Printer (Lo3DP), a reconfigurable robotic microfluidic platform that repurposes the motion control, thermal regulation, and open-source programmability of consumer 3D printers to deliver laboratory-grade automation at a hardware cost below $500. By replacing the extruder with a multifunctional magnetic toolhead, Lo3DP actuates ferrofluid droplets with sub-100 µm positioning accuracy and programmable volume control spanning 0.5–25 µL, enabling the full repertoire of digital microfluidic operations within inexpensive laser-cut chips (<$3). We quantitatively validate system performance through long-term droplet transport exceeding 40,000 s without degradation, automated serial dilutions with high linearity (R² ≥ 0.99), and fully automated colorimetric nucleic acid amplification assays whose results match conventional benchtop workflows under tightly regulated isothermal conditions (CV ≈ 0.3%). By harnessing the global economies of scale of consumer 3D printers, Lo3DP provides accurate, reproducible, and programmable assay automation at a fraction of the cost and footprint of conventional systems, establishing a practical path toward desktop-scale, democratized laboratory robotics.

## I. Introduction

Automation has revolutionized many sectors of science and engineering, yet life science research remains mostly manual, labor-intensive, and cost-prohibitive to scale. Robotic liquid-handling systems have accelerated industrial drug discovery^1,2^, biomedical research^3^, diagnostics^4^, and biomanufacturing^5^ by enhancing throughput and reproducibility while simultaneously reducing errors^6^. However, these platforms are refrigerator-sized and typically priced at tens of thousands to hundreds of thousands of dollars. They are optimized for centralized facilities^7^ and limited to larger liquid volumes compatible with standardized multi-well plates. As a result, high-performance lab automation remains inaccessible to most academic researchers, small biotechnology labs, and other resource-limited settings^8^. Achieving a more decentralized and affordable form of experimental automation requires reimagining not only the scale of instruments but also the fundamental architecture for conducting liquid bioassays.

Digital microfluidics (DMF) has been proposed as a promising route to miniaturized laboratory automation. By actuating microscale droplets via electrowetting-on-dielectric (EWOD)^9^ or electret-induced polarization on droplets (EPD)^10^, DMF platforms can execute complex bioassay workflows with sub-microliter to microliter precision. More recently, PCB-based ferrobotic systems^11,12^ have extended this concept by magnetically manipulating droplets over an array of addressable electromagnetic coils. Despite their technical sophistication, current DMF and ferrobotic systems remain difficult to deploy broadly. Their multilayer fabrication processes, high-voltage or complex PCB control electronics, weak magnetic actuation strength, and lack of integrated sample introduction limit scalability, robustness, and usability^11,13,14^. These constraints underscore the need for robust, self-contained automation platforms that achieve microscale precision without relying on expensive or complex infrastructure.

The rapid advancement of consumer-grade 3D printing offers an unexpected but scalable solution to fill this gap. Modern 3D printers incorporate capabilities similar to those of laboratory robots, including high-precision motion control, temperature regulation, and open-source programmability via G-Code. Because these printers are mass-produced for global consumer markets, their economies of scale reduce hardware costs by several orders of magnitude^15,16^. Systems costing <$400 rival the positional accuracy of entry-level liquid handlers that cost ∼ $15,000^15^. Recent repurposing efforts demonstrate this potential (**Table S1**): modified printers such as the Creality Ender-3 have performed automated histology staining for <$215^17^, the D-Bot Core-XY has conducted remote fluorescent imaging^18^, and the Printrbot Simple has executed rapid nucleic-acid purification^19^. These proof-of-concept demonstrations reveal that consumer 3D printers can match, and sometimes even surpass, specialized laboratory robots in precision, modularity, and reproducibility. Therefore, we sought to integrate these advantages of 3D printers with DMF droplet control into a unified, programmable platform for versatile, low-cost laboratory automation.

Here, we report ***Lab-on-a-3D-Printer*** (**Lo3DP**) (**Fig. 1 *right***) to miniaturize and decentralize the tools for laboratory automation. It utilizes a modified, consumer-grade 3D printer with its extruder replaced by a multifunctional manipulation head to perform routine lab operations. Compared to conventional robotic liquid handlers (**Fig. 1 *left***) that rely on well-plate architectures and an expensive gantry system, the Lo3DP reduces both the physical footprint and the reagent consumption by an order of magnitude (**Table S2**). The Lo3DP system achieves this miniaturization by using ferrofluid-infused magnetic droplets (ferrodroplets) that can be precisely transported and manipulated with stacked magnets on its manipulation head. The Lo3DP also inherits the high positioning accuracy of the base 3D printer, resulting from high-precision motors that encode accurate localization for polymer extrusion and printing. By sending G-code^20^ commands to move the stacked magnets, droplet transportation can be achieved. This magnetically driven approach replaces conventional pipetting mechanisms, enabling droplet-based operations within a microfluidic chip. Thermal control of the print bed can also be used to control the reaction temperatures on the Lo3DP platform.

**Fig. 1.**
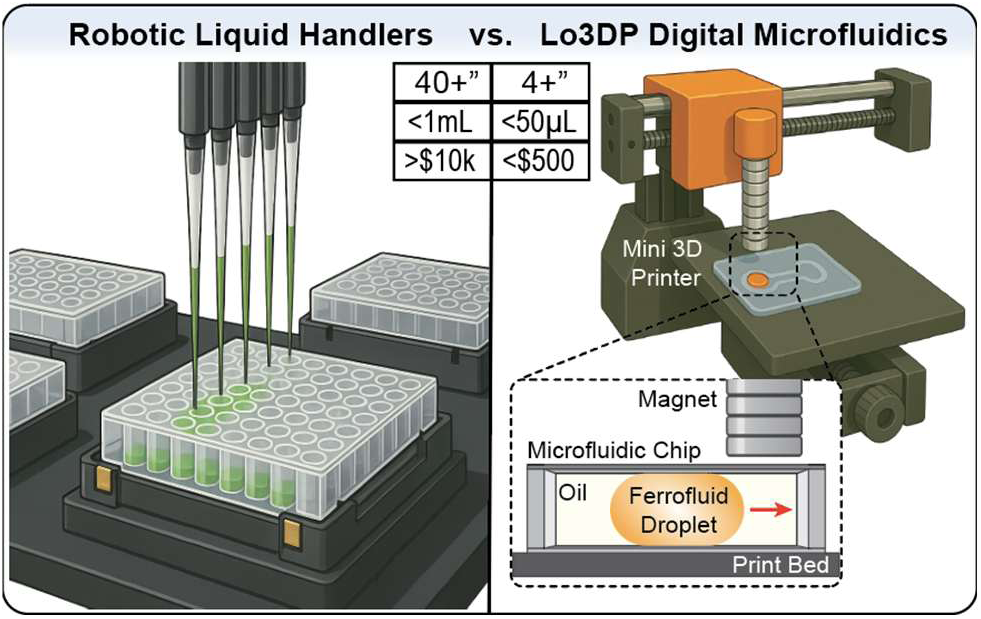
Miniaturization of laboratory automation with Lo3DP. Schematics showing a robotic liquid handler with standard well plates and Lab on a 3D-Printer (Lo3DP), comparing instrument size and cost, as well as reagent volume. The inset shows the mechanism of liquid transport and manipulation with a mini 3D printer with a magnetic manipulation head to attract ferrofluid-infused droplets (ferrodroplets).

Everyday laboratory tasks, such as liquid introduction, mixing, serial dilution, incubation, magnetic bead separation, and optical detection, were implemented on a single Lo3DP platform. Each operation is programmed through G-Code instructions that direct ferrodroplets within assay-specific microfluidic chips. Implementing these operations can lead to streamlined, automated execution of multistep experimental protocols on a benchtop-sized system while dramatically lowering cost and reagent use compared to conventional robotic laboratory automation (**Table S2**). More broadly, the Lo3DP demonstrates that the precision mechatronics and thermal control embedded in mass-produced consumer devices can be repurposed to deliver true laboratory-grade automation. As these repurposable consumer-robotics platforms proliferate, we anticipate a ripple effect across the field, shifting the center of gravity of laboratory automation from highly engineered, single-purpose instruments toward adaptable and widely distributed robotic systems that democratize experimental precision and expand participation in cutting-edge biological research.

## II. Results

### II. 1) System Infrastructure and Automated Microfluidic Operations with the Lo3DP

To achieve high versatility within the compact Lo3DP framework, we constructed a fully integrated hardware and software infrastructure capable of performing programmable, droplet-based bioassays (**Fig. 2 and Fig. S1**). The Lo3DP system consists of components on the print bed, as well as on the frame and external areas (**Fig. 2A**). On the print bed, tube & tip racks and a pipettor tip ejector are dedicated to liquid sample preparation and introduction. The microfluidic chip, filled with an immiscible oil environment surrounding ferrodroplets, is engineered to perform digital microfluidic bioassays. The frame and external components include a touchscreen for user-Lo3DP interaction, a liquid handling module, and a manipulation head that enables numerous functions, such as liquid introduction, ferrodroplet transportation and merging, and optical detection.

**Fig. 2.**
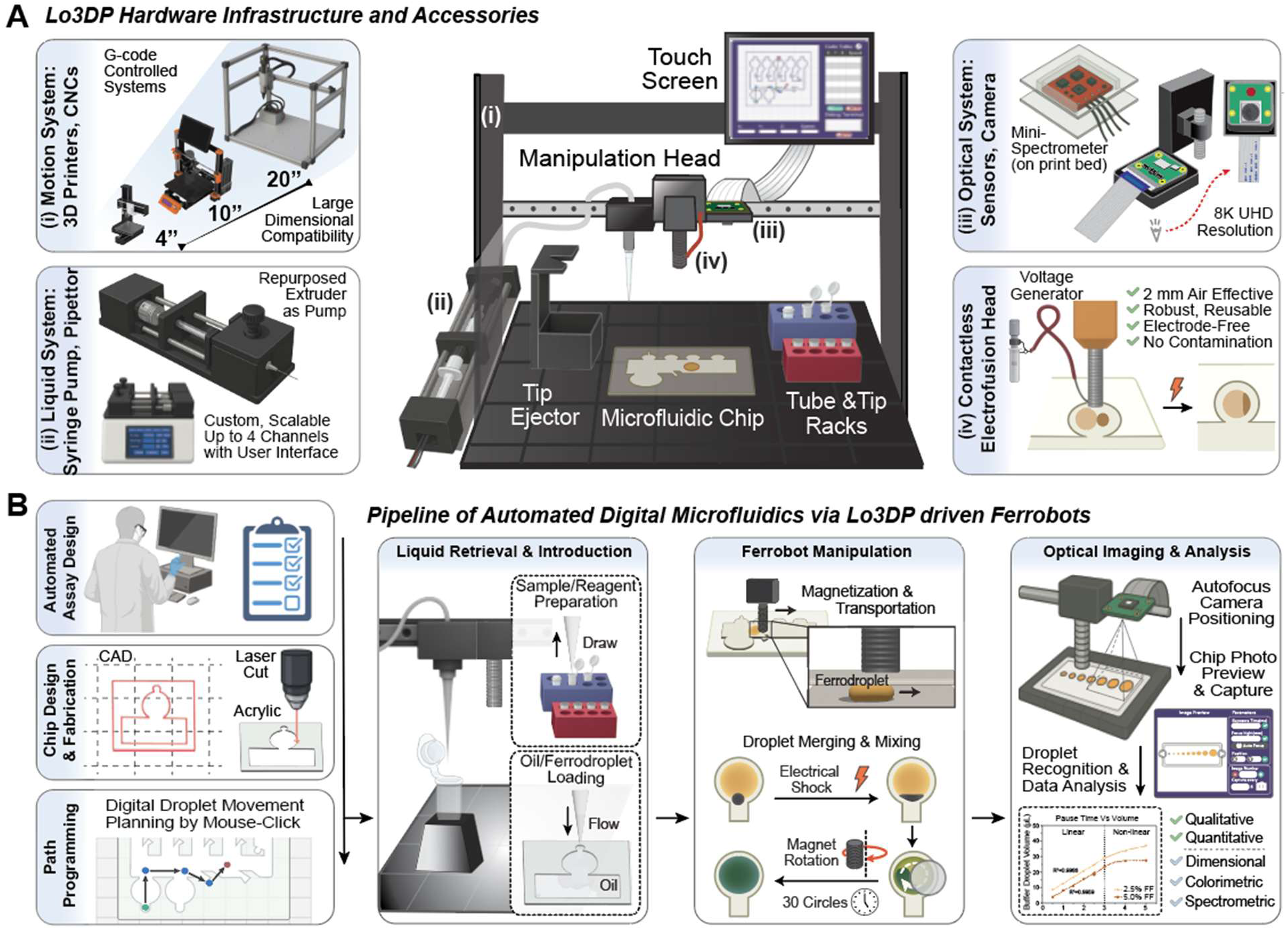
Infrastructure and automated microfluidics with Lo3DP. **(A)** Illustrations showing the design and setup of the Lo3DP: components on the print bed include a microfluidic chip for droplet-based bioassays, tube & tip racks for holding tubes and pipette tips, and a tip ejector for pipette tip removal; other components include (i) an x-y-z motion system: a custom G-Code controlled 3D motion gantry from 4“ to 20” (leveraging 3D printers, CNCs, and liquid handlers), (ii) a liquid control system: a custom, scalable syringe pump (repurposed from the 3D printer filament extruder) connecting to a pipettor mounted on the manipulation head (MH), (iii) an optical system: a mini-spectrophotometric sensor and a high-resolution Raspberry Pi camera, and (iv) an electrofusion head: a voltage generator (connected to stacked magnets) delivering a transient electrical shock without a contact to disrupt the surfactant shell and allow ferrodroplet merging. **(B)** A streamlined pipeline from bioassay design to data analysis via the Lo3DP: 1) Researchers design the automated procedure and corresponding microfluidic chip. Then, import the CAD design of the microfluidic chip into the user interface and click on the grid to form a movement path for the magnet. 2) The pipettor on the MH collects oil and fills the microfluidic chip. Then, it prepares and draws aqueous ferrofluid-infused samples and reagents from tubes and injects them into the microfluidic chip to form ferrodroplets. And then the pipette tip is ejected. 3) The stacked magnets on the MH perform a sequence of programmed ferrodroplet operations, like transporting, dispensing, merging, and mixing. 4) Droplet imaging and measurement are performed using an autofocus camera or a spectrophotometric sensor, and subsequent recognition and quantitative analysis are automatically conducted.

The external components are crucial for creating a modular infrastructure for miniaturized lab automation, as shown in **Fig. 2A**. Due to G-Code’s universal compatibility, our Lo3DP concept is generalizable to broader 3D gantry systems (**Fig. 2A (i)**), like low-cost consumer-grade printers (∼4’’), standard 3D printers (∼10’’), and liquid handling robots (∼20’’), plus other CNC motion systems. A repurposed 3D-printer extruder-based liquid handling system (**Fig. 2A (ii)**) can be expanded to 4 channels and controlled by G-Code commands, enabling easy programming and integration of liquid-transfer tasks. There are three major components mounted on the manipulation head: a magnetic/electrofusion head for droplet transportation and contactless merging (**Fig. 2A (iv)**), a pipette-tip adaptor, and an optical module (*i.e.*, camera and sensors) (**Fig. 2A (iii)**). The user can enter coordinates or create motion paths on the touchscreen interface, and G-Code will be generated to control the precise localization of various functional components.

The Lo3DP workflow outlines the operational sequence from experimental design to data analysis. It can be divided into two major parts: design (**Fig. 2B *vertical***) and execution (**Fig. 2B *horizontal***). The design process includes the creation of bioassay procedures, microfluidic chip design & rapid laser-cut fabrication, and spatial motion path programming through a graphical user interface. Next, the execution process comprises liquid retrieval & introduction, ferrodroplet manipulation, and optical imaging & analysis. In detail, the pipettor on the manipulation head collects and dispenses oil and ferrofluid-infused aqueous reagents to form ferrodroplets within the microfluidic chip. The system subsequently executes a programmed series of ferrodroplet operations, such as transportation, dispensing, merging, and mixing. After that, the optical module automatically performs image capture or spectrophotometric measurement, followed by algorithmic data recognition and quantitative analysis.

### II. 2) Characterization and Quantitative Performance of Ferrodroplet Manipulation

Microfluidic automation is the key to miniaturizing the experimental footprint and reaction volumes. We conducted a series of quantitative experiments to evaluate the Lo3DP’s capabilities in ferrodroplet transportation, dispensing, mixing, and magnetic bead (MB) separation (**Fig. 3**). These modular operations are fundamental to matching standard lab procedures and automating assays (**Fig. 1B**).

**Fig. 3.**
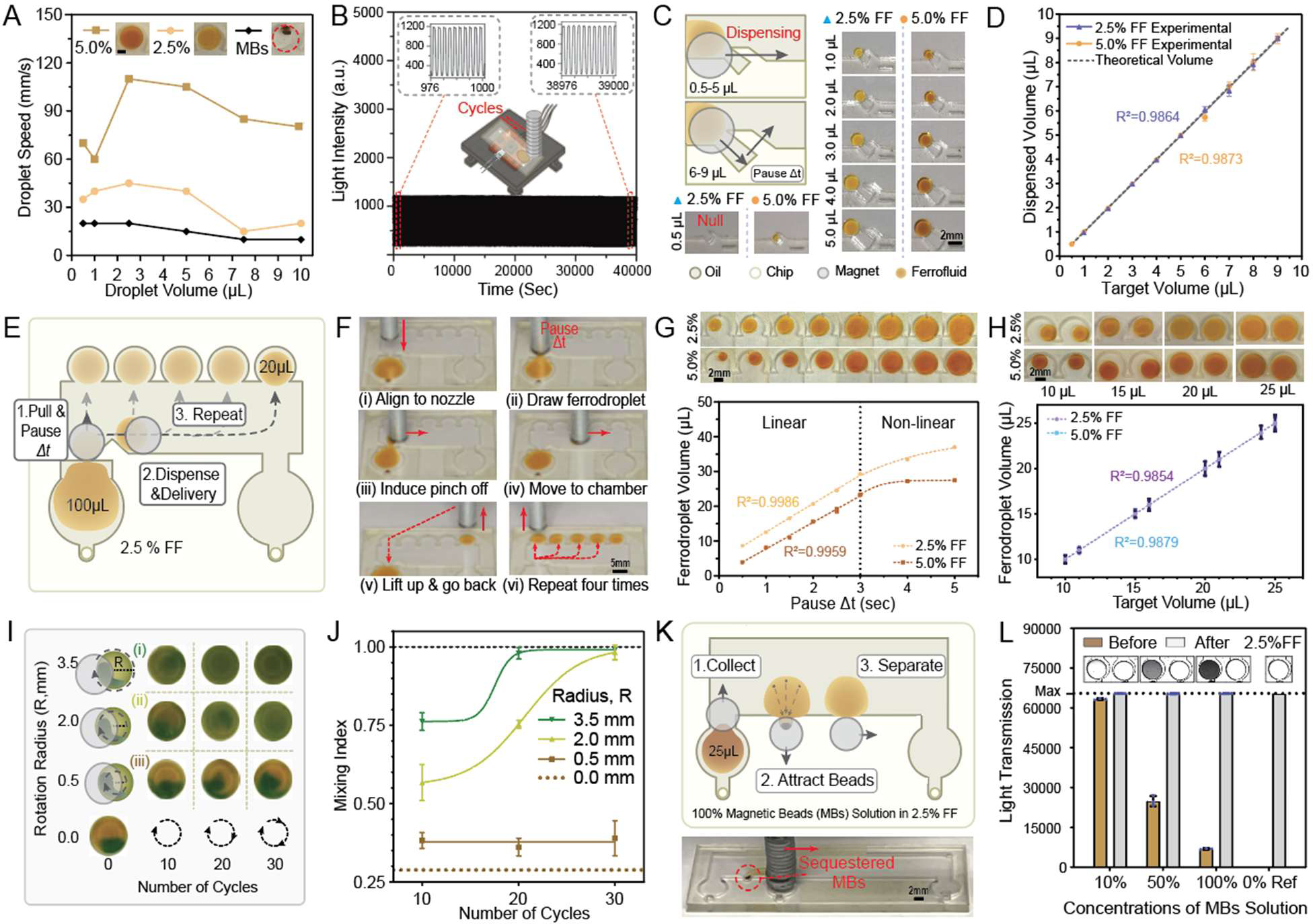
Characterization of ferrodroplet manipulation with Lo3DP. **(A)** Characterization of the transportation speed of droplets infused with ferrofluids (FF, 2.5% or 5.0%, uniformly distributed under a magnetic field) or magnetic beads (MBs) (clustered). **(B)** Robust droplet transportation was verified by moving a 2.5% ferrodroplet across a spectrophotometric sensor, occluding it, and then moving back to the origin, and repeating this transportation over 40,000 sec. The sensor detects the repetitive patterns as shown in the insets. The illustration shows the test setup on the print bed. **(C)** Characterization of the dispenser for tiny volume aliquoting. The magnet moves straight across the dispenser to create 0.5-5 μL daughter droplets, or moves diagonally over, pauses, and moves out of the dispenser to create 6-9 μL daughter droplets. **(D)** The volume accuracy of dispensed droplets with 2.5% or 5.0% FF is plotted, showing high correlation (2.5%, R^2^ = 0.99; 5%, R^2^ = 0.99). **(E)** An illustration of the droplet generator layout and major steps of creating multiple daughter droplets (Steps explained in **F**). **(F)** Images of the pipeline to generate five 20 μL daughter droplets: (i) Align the magnet to the nozzle and slowly move down to magnetically attract the parent ferrodroplet (100 μL). (ii) A part of the ferrodroplet flows out of the nozzle (width of 1.0 mm), and the pause time (Δt) controls the volume of a daughter droplet. (iii) Quickly move the magnet to the right to pinch off the daughter droplet. (iv) Move the daughter droplet to the storage chamber. (v) Lift the magnet and go back to the parent droplet position. (vi) Repeat four times until the parent droplet is fully consumed. **(G)** Relationships between pause time **Δt** (sec) and the generated ferrodroplet volume (μL) are plotted. Before the daughter droplet size reaches the size of the magnet, a strong linear relationship exists (2.5%, R^2^ = 0.99; 5.0%, R^2^ = 0.99). **(H)** The volume accuracy was plotted, showing the generated daughter droplets of different target volumes between 10 and 25 μL (2.5%, R^2^ = 0.99; 5.0%, R^2^ = 0.99). **(I)** Characterization of magnetic mixing of ferrodroplet contents. A 2 μL dye sample was electrofused with 8 μL buffer droplets and magnetically mixed by using different combinations of the number of rotation cycles and radii. Mixing images are shown from 10 to 30 cycles of rotations with a radius of 0 to 3.5 mm. **(J)** The Mixing Index (dye-colored area ratio) is plotted for different numbers of rotation cycles and radii. **(K)** An illustration of the MBs separator layout and major steps of bead capture. The stacked magnets align with the MBs-infused ferrodroplet (Step 1), then moves the droplet against a semicircular chamber (radius of ∼0.5 mm) while the MBs are clustered and immobilized (Step 2). The magnet then extracts the ferrodroplet from the MBs, moving the droplet to the right (Step 3). The sequestered MBs are indicated in the bottom image by a red dashed circle. **(L)** Separation efficiency was quantified by comparing light transmission before and after separation. After separation, all the groups (10%, 50%, and 100% MBs solutions) achieved the same level of transparency compared to the 0% reference group. Obvious color changes can be observed before and after the separation, as shown in the insets for the 50% and 100% MBs solutions.

Droplet transportation speed is one contributing factor to the operating time of the Lo3DP automation technology. Ferrodroplet transportation speeds were measured by setting a specific feed rate (F value) in G-Code commands. Maximum speeds that the ferrodroplet could follow with a 4.5 cm displacement of the stacked magnets were recorded in **Table S4** and plotted in **Fig. 3A**. Droplets with ferrofluid at concentrations of 2.5% and 5.0% maintained high mobilities, comparable to previously reported speeds^11^ where even higher ferrofluid concentrations (*e.g.*, 7.5%, 10.0%, and 12.5%) were required. In contrast, MB-laden droplets exhibited clustering and slower transportation, further revealing the superior homogeneity and magnetic responsiveness of ferrodroplets for digital microfluidic operations. Both ferrodroplets (with lower ferrofluid concentrations) (**Note S1**) and MB-laden droplets achieved high levels of optical transparency.

Robust, repetitive ferrodroplet operations are essential for practical applications and a broader adoption of the Lo3DP concept. Repeated transportation of 2.5% ferrodroplets was demonstrated in **Fig. 3B**. Specifically, the stacked magnets of the manipulation head moved the ferrodroplet across a spectrophotometric sensor, repeatedly occluding it and returning to the origin at a rate of 0.4 cycles per second for 40,000 seconds. The optical readout curve showed light intensity (a.u.) over these repeat cycles. The curves of two representative time periods, as shown in **Fig. 3B *Inset*** 976∼1,000s and ***Inset*** 38,976∼39,000s, exhibited identical motion patterns, validating long-term transportation repeatability.

Dispensing reagents and samples into smaller fractions is fundamental for life science research. As shown in **Fig. 3C-H**, we achieved high-precision dispensing and droplet generation with a well-suited volume coverage, 0.5∼25 μL, by utilizing two types of droplet manipulation structures, including hook-based dispensers (**Movie S1**) and nozzle-based droplet generators (**Movie S3**). The Lo3DP moves the stacked magnets to drag a parent ferrodroplet across hook-based structures (**Fig. 3C**) in the microfluidic chip and generate 0.5∼9 μL daughter ferrodroplets. **Fig. 3D** shows that the dispensed volumes matched well with the target volumes programmed through the hook geometry for two different ferrofluid concentrations (2.5% ferrofluid (FF): R² = 0.99; 5.0% FF: R² = 0.99). The deterministic, consistent dispensing capability across a wide range of FF concentrations (2.5% - 15%) has been validated (**Note S2** and **Movie S2**).

Once the target droplet volume exceeds 9 μL, it becomes suboptimal to operate using the stacked magnets with a 3 mm diameter. One workaround is to increase the diameter of the stacked magnets, which introduces a stronger magnetic field and a larger cross-sectional area. However, using larger stacked magnets to manipulate a ferrodroplet can result in a lower positioning accuracy and a higher risk of attracting non-targeted ferrodroplets. Thus, we developed a nozzle-based droplet generator that can dispense a larger amount of liquid from 10 to 25 μL (**Fig. 3E** and **Fig. S2**). By positioning the stacked magnets over the nozzle neck (**Fig. 3F (i)**), a part of the fluid from the parent ferrodroplet flows out from the nozzle (**Fig. 3F (ii)**). The period during which the stacked magnets are overhanging is defined as pause time **Δt** (sec). By precisely controlling the **Δt**, specified amounts of liquid flow from the nozzle and are pinched off (**Fig. 3F (iii)**). This droplet generation is repeated several times to create multiple copies for parallel reactions (**Fig. 3F (vi)**). The individual steps of ferrodroplet manipulation to create many aliquots are detailed in the caption of **Fig. 3F**.

We investigated the correlation between **Δt** and the resulting volumes of nozzle-generated ferrodroplets. **Fig. 3G** showed that the relationships exhibited excellent linearity (R² = 0.99) for both 2.5% FF and 5.0% FF before **Δt** reached ∼3 sec, where the volumes of fluid flowing out of the nozzle were proportional to **Δt**. When the sizes of daughter ferrodroplets reached the size of the stacked magnets, the volumes tended to be saturated, and relationships between **Δt** and volume transitioned to be non-linear. Based on the quantitative relationship acquired above, our nozzle-based droplet generators achieved target volumes ranging from 10 to 25 μL (**Fig. 3H**). The operations with 2.5% and 5.0% FF both showed high linearity (R² = 0.99) across eight target volumes, providing a foundation for accurate downstream bioassays.

To achieve bioassay reproducibility, reagent mixing is a key step that ensures a uniform distribution of assay contents in a liquid environment. We conceptualized a “magnetic vortexing” method that involves rotation of the stacked magnets over a heterogeneous ferrodroplet in a mixing chamber. Magnetic mixing efficiency was quantified by measuring *Mixing Indices* of mixed, dye-colored ferrodroplets (**Fig. 3I**). Controlled rotational motions of the stacked magnets stirred the inner contents of an unmixed ferrodroplet and enabled uniform color homogenization within 30 cycles with a rotation radius of 3.5 mm and a motion speed at 0.5 cm/s (**Fig. 3J**). These results demonstrate the Lo3DP’s capability to efficiently homogenize droplet contents through programmable magnetic stirring, without physical agitation or contamination.

Magnetic beads (MBs) are commonly used in solid-phase bioassays to selectively enrich analytes, usually equipped with functional groups on their surfaces for specific binding of target molecules. Separating MBs is an important step for washing out non-target molecules or concentrating analytes of interest. We validated this sequestering capacity using the Lo3DP system (**Fig. 3K**). By moving the stacked magnets along predefined paths, MBs were successfully immobilized against a semicircular chamber wall while the surrounding ferrofluid was extracted, effectively isolating the beads (**Movie S4**). As shown in **Fig. 3L**, optical transmission analysis and microscopic images revealed that post-separation transparency levels in 10%, 50%, and 100% MB solutions were identical to those of the 0% reference control, confirming complete removal of magnetic particles.

In short, the above results validated that the Lo3DP platform can perform modular microfluidic manipulation operations needed for various bioassays in a robust and repeatable manner.

### II. 3) Automated Serial Dilution with Lo3DP

To evaluate the Lo3DP’s capability for automating workflows consisting of a multi-step sequence of the demonstrated liquid handling tasks (*e.g.*, droplet transportation, dispensing, and mixing), the Lo3DP was programmed to automate serial dilutions using ferrodroplets (**Fig. 4 and Movie S5**). **Fig. 4A** shows graphical illustrations of the process—a sequence of magnetic dispensing, transport, electrofusion, and mixing steps digitally controlled by the Lo3DP without human intervention. In detail, the two major steps are i) Buffer Droplet Dispensing: collecting the parent buffer droplets from reservoirs, dispensing them into daughter droplets with designated volumes (*e.g*., 8 μL for 5× serial dilution), and distributing daughter droplets to the dilution chambers located on the top row; ii) Dye Sample Dilution: dispensing the sample into a daughter droplet (*e.g*., 2 μL for 5×) and transporting it to the proceeding dilution chamber for contactless electrofusion and mixing by magnetic stirring. This process iterates several times until the final dilution in the last chamber is completed. Individual ferrodroplet manipulation steps are detailed in **Note S3** and the caption of **Fig. 4A**.

**Fig. 4.**
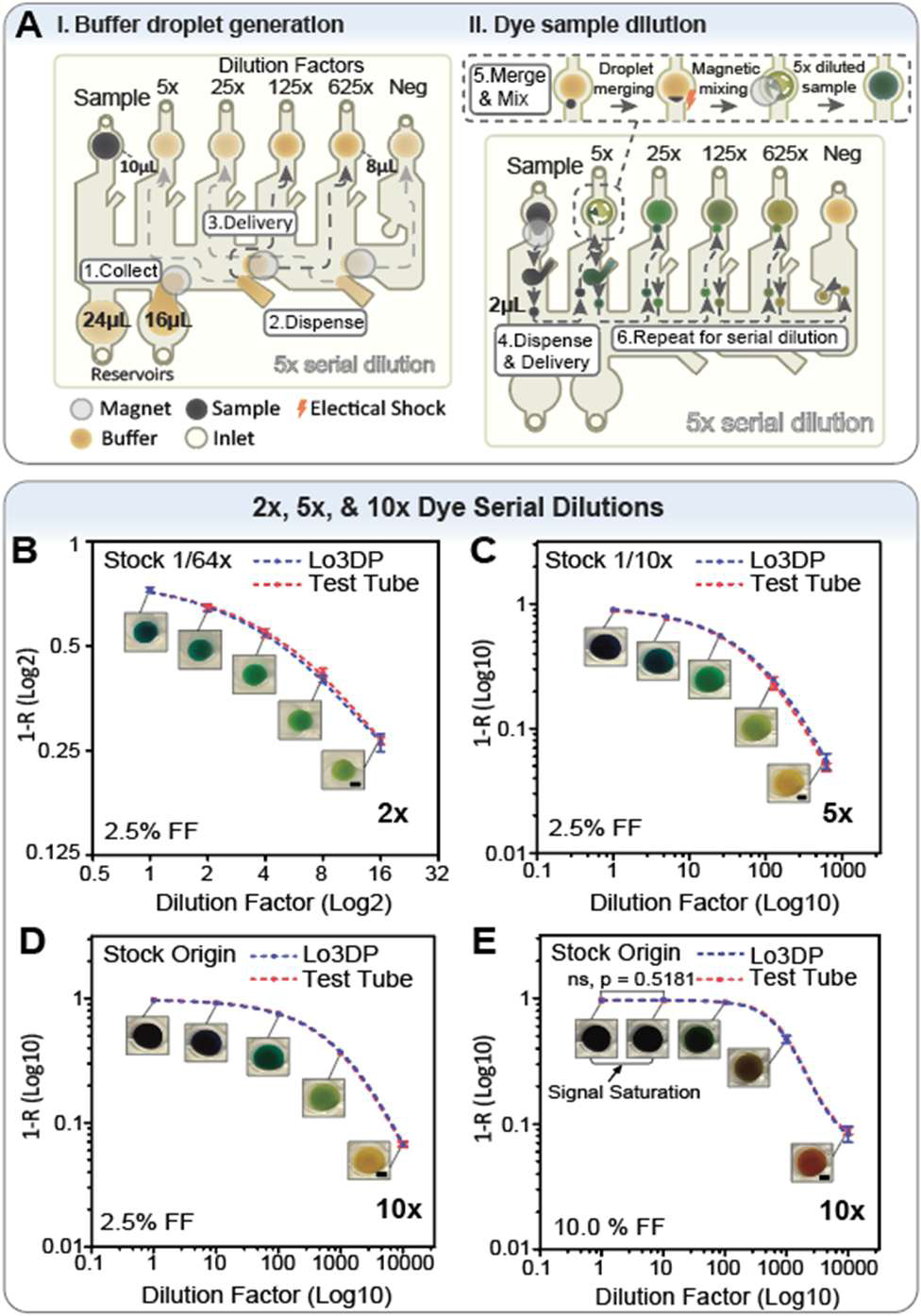
Automated serial dilutions with Lo3DP. **(A)** A schematic workflow illustration of a 5× serial dilution, including buffer droplet generation and dye sample dilution: (i) the buffer droplets were formed by collecting the ferrofluid (FF)-infused parent buffer drops from the buffer reservoirs (Step 1), dispensing them into individual daughter drops (Step 2), and then moving the daughter droplets up to the dilution chambers (Step 3). (ii) The dye samples, also infused with ferrofluid, were serially diluted by dispensing the sample and delivering it to the closest dilution chamber with a daughter buffer drop (Step 4). The dispensed sample and the daughter buffer drop are merged using contactless electrofusion, and the merged droplet is rotated magnetically to mix its contents (Step 5). Finally, this process was repeated several times across the sequential dilution chambers (Step 6). Graphs showing the normalized light intensity (1-*R ratio*)* *vs* dilution factor* for 2× **(B)**, 5× **(C)**, and 10× **(D)** serial dilutions with the Lo3DP using 2.5% FF droplets. The decreasing colorimetric signal with dilution matched well between the Lo3DP and the manual test tube methods. **(E)** A graph showing the normalized light intensity (1-*R ratio*)* *vs* the dilution factor* for 10× serial dilution with Lo3DP using 10% FF droplets, showing signal saturation (arrowed on graph) that obscures the colorimetric readouts (Color signals were not significantly different, p = 0.5181, when diluting the sample 10 times). The curves of colorimetric signals matched well between the Lo3DP and manual test tube methods. *Log2 scale for 2× and Log10 scale for 5×/10× serial dilutions.

Quantitative testing evaluated the accuracy of serial dilutions with the Lo3DP. Several dilution experiments (*i.e.*, 2×, 5×, and 10×) were conducted to verify the capabilities and robustness of this system. In detail, the Lo3DP system was programmed to serially dilute a dye sample from a starting concentration (*e.g*., 1/10× stock for 5×) to concentrations several orders of magnitude lower (*e.g*., dilution factors after each step of 5, 25, 125, and 625 for 5×), followed by image acquisition and colorimetric quantification.

As shown in the **Figs. 4B-C**, colorimetric changes from darker to lighter can be observed. For example, when the dilution factor reached 10,000 during the 10× serial dilution, the original light-brown color of the ferrodroplet became observable. We also conducted standard dilutions with test tubes, pipettors, and a vortexer to measure comparative colorimetric changes to be plotted alongside Lo3DP measurements. Results showed that the normalized light intensity (1-*R ratio*) of the two methods matched well (2×: R² *=* 0.99; 5×: R² *=* 0.99; 10×: R² *=* 1.00) with minimal deviations across replicate trials.

An advantage of the Lo3DP is the high magnetic strength of its stacked magnets, allowing for digital microfluidic operations with minimal concentrations of infused ferrofluid (FF). By comparing the colorimetric signal curves in **Fig. 4D** and **Fig. 4E**, an obvious signal saturation (p = 0.52 > 0.05, indicating no significant colorimetric changes after diluting the original stock sample 10 times) can be observed with 10% ferrodroplets utilized. The droplet images shown below the saturated signal curve remain uniformly dark, showing that excessive iron oxide nanoparticles can obscure optical signals. Further validation of this optical advantage is detailed in **Note S1** and **Fig. S3**. Despite the aforementioned signal saturation, the light intensities of Lo3DP and Test Tube groups still matched well at 10× with 10% FF (R^2^ *=* 0.99), showing robust performance across diverse fluid compositions.

### II. 4) Automated Colorimetric LAMP with Lo3DP

To assess the applicability of the Lo3DP for molecular diagnostics, we implemented a fully automated colorimetric loop-mediated isothermal amplification (**LAMP**) workflow for BRCA1 primer set validation in breast cancer diagnostics (**Fig. 5**). The assay leveraged the Lo3DP’s integrated fluid handling, droplet electrofusion, thermal regulation, and imaging capabilities to execute all steps of a LAMP reaction without manual operation. As illustrated in **Fig. 5A**, after loading the initial reagents (*e.g.*, LAMP primer sets (i)-(v), positive DNA (+ DNA), negative control, and LAMP master mix), the workflow began with the dispensing of primer set droplets, followed by + DNA & negative control droplets. These reagents were merged into LAMP mix droplets located in the top chambers (**Movie S6**). Then, the print bed of the Lo3DP system was heated up to 65 ℃ for 30 minutes of isothermal amplification, followed by image capture and analysis.

**Fig. 5.**
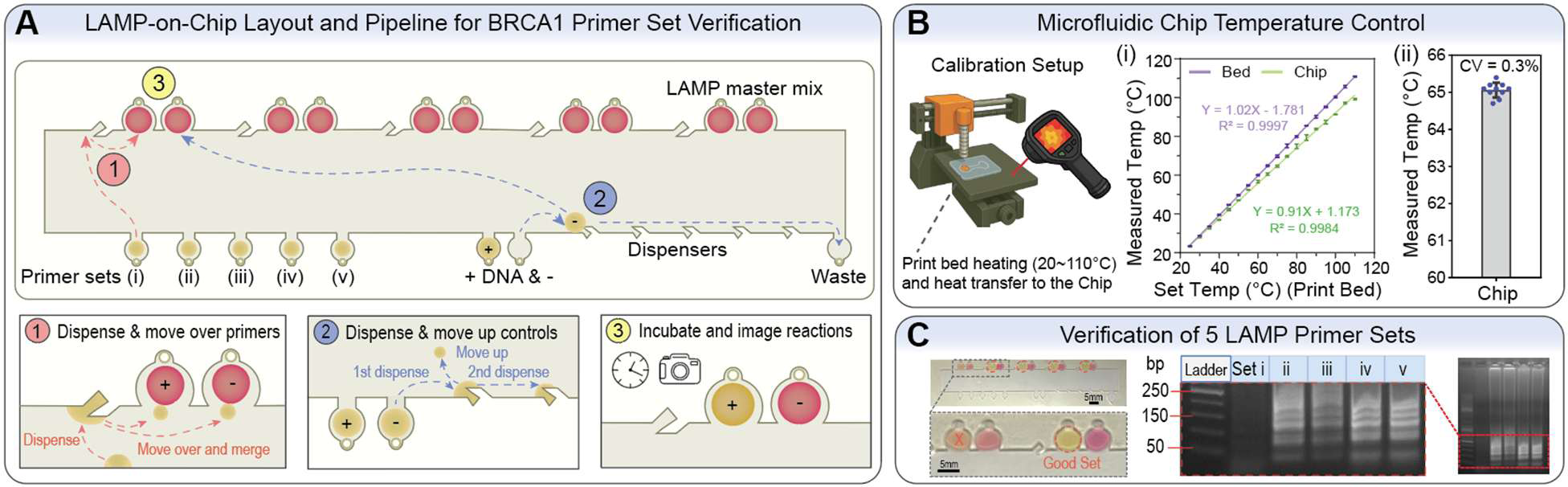
Automated Colorimetric LAMP-on-Chip with Lo3DP. **(A)** A schematic of the LAMP assay workflow for automated in-parallel testing of primer sets. The initial positions of reagents such as LAMP primer sets (i)-(v), positive DNA and negative control (+ DNA & -), and LAMP master mix are shown. The workflow for verifying BRCA1 LAMP primer sets involves several steps following the loading of oil and all reagents: 1) Dispense and transport primer sets: the primer set (i) is moved to the dispenser adjacent to the LAMP master mix to split the parent droplet into two droplets, followed by sequential addition to the reaction mix droplets. This process is repeated for primer sets (ii)-(v). 2) Dispense and transport + DNA & - samples to merge with the LAMP master mix: move the negative control parent droplet to the dispensers to form five daughter ferrodroplets, which are then merged with five different LAMP master mix droplets to form reaction mix droplets. Then, this process is applied to the DNA + parent droplet. 3) Incubate and capture reaction images: perform *in situ* heating to 65℃, incubate for 30 minutes, and acquire endpoint colorimetric images of the LAMP reactions. **(B)** Microfluidic chip temperature control and calibration setup: the illustration depicts the calibration process in which the 3D printer print bed is heated from 20℃ to 110℃ while the thermal camera measures the temperature of both the print bed and microfluidic chip. A strong linear correlation is observed between the set temperature and chip temperature (R^2^ = 1.00), with temperature stability at 65℃ measured over ten trials (CV of 0.3%). **(C)** Validation of primer sets for BRCA1 breast cancer gene detection by colorimetric LAMP on the Lo3DP. Successful amplification of the target gene induces a pH change, resulting in a clear color transition from pink to yellow. The positive droplet from primer set (i) remained pinkish, indicating insufficient amplification and a suboptimal primer set. In contrast, the positive droplets from the other primer sets (ii)-(v) displayed yellow coloration after incubation, indicating efficient amplification and effective primer sets, which matched well with gel electrophoresis, revealing distinct amplicon bands in sets (ii)-(v).

Thermal regulation and calibration of the microfluidic chip were conducted as shown in **Fig. 5B**. The Lo3DP print bed provided uniform and accurate heating from ∼25 to 110 °C. After heat transfer from the print bed to the microfluidic chips, there was an excellent linear correlation between the programmed print bed temperature and the measured microfluidic chip temperature (Y = 0.91X + 1.173; R² = 1.00). Based on the equation, we achieved temperature control on a microfluidic chip over a programmable range from ∼25 to 100 °C. Temperature stability during isothermal incubation at 65 °C was exceptional, with a coefficient of variation (CV) of only 0.3 % over ten independent trials. This result confirmed that the 3D printer’s heated bed can be directly harnessed to maintain precise thermal conditions for bioassays.

As shown in **Fig. 5C**, we demonstrated an on-chip colorimetric LAMP assay that validated efficient primer sets for breast cancer detection with the BRCA1 target gene. After incubation, reactions with an efficient primer set produced a visible color change from pink to yellow due to a decrease in pH associated with efficient DNA amplification, while negative controls remained pink. Among the five primer sets tested, set (i) showed minimal color transition, indicating inefficient amplification, whereas sets (ii)–(v) yielded clear color changes to yellow, indicating successful amplification. The Lo3DP-based colorimetric results closely matched gel electrophoresis outcomes (**Fig. S4**) for bulk assays, which confirmed amplicon formation only in sets (ii)–(v). These findings demonstrated that Lo3DP can reliably perform automated, in parallel nucleic acid amplification assays with integrated colorimetric detection.

## III. Discussion

While large-scale robotic systems and humanoid robots have been proposed for laboratory automation^6,21^, costs and complexity have hindered their widespread adoption. Large-scale robots occupy entire benches or rooms to run single-purpose bioassays, requiring specialized personnel and upfront investment beyond the reach of most academic laboratories^22^. General-purpose humanoid robots are constrained by their anthropomorphic design that introduces complexity in gait control, wrist articulation, and collision avoidance^23^. Their kilogram-scale actuators and safety enclosures increase cost and energy use, yet still fall short of the submillimeter precision required for droplet-based assays. In contrast, the Lo3DP is purpose-optimized for laboratory tasks, directly manipulating microliter-scale ferrodroplets with sub-100 µm repeatability using modified, low-cost 3D printers that fit entirely within a biosafety cabinet. By leveraging open-source electronics and modular hardware, the Lo3DP delivers a compact, scalable, and efficient alternative that surpasses both large-scale automation and humanoids in precision and accessibility.

Owing to the economies of scale associated with the widespread deployment of 3D printers^24^, the Lo3DP achieves an exceptionally low hardware cost compared to standard liquid handlers and conventional microfluidic systems, often below $500 (**Table S2**). Existing low-cost custom liquid handling systems still have drawbacks in precision and throughput (**Table S3**). Lab-on-a-Chip technologies have achieved remarkable miniaturization and precision of reaction volumes^25^. However, their commercial implementation and widespread use have been limited by their reliance on bulky and costly instruments^25^, such as precision pumps for fluid control and high-voltage electronic controllers for EWOD actuation. The Lo3DP balances miniaturization and functionality, providing digital microfluidic control within a compact, self-contained platform that integrates droplet manipulation, thermal regulation, and real-time imaging. Moreover, by leveraging magnetic droplet actuation, Lo3DP enables programmable, non-contact liquid handling without mechanical wear or intricate circuitry, enhancing reliability and ease of use.

The Lo3DP shares the fundamental principle of magnetic droplet actuation with swarm ferrobotics based on an addressable PCB coil array^12^. The swarm ferrobotic system manages multiple mobile magnets atop a PCB using sophisticated path-planning algorithms; however, neighboring magnets could attract each other, collide, or unintentionally attract non-target ferrodroplets. In contrast, the Lo3DP trades off parallelization with robustness, employing a single column of stacked magnets, enabling smooth droplet localization through point-to-point, collision-free movement control. The Lo3DP system also integrates sample preparation and loading using a pipetting module, which retrieves reagents from tubes and introduces them into a microfluidic chip. This capability is absent in swarm ferrobotics, which typically operates on preloaded droplets. The Lo3DP’s stacked magnets (>10 mm) produce magnetic fields much stronger than those of a single-layer ferrobotic magnet, enabling more rapid, reliable transportation of low-concentration ferrodroplets with high optical transparency. This combination yields robust actuation, quality imaging, and compatibility with diverse bioassays. While swarm ferrobotics excels at parallel droplet-moving tasks, the simple, integrated Lo3DP ecosystem is more practical and efficient for most lab automation processes, where a majority of assay time is spent in incubation (>10 mins) rather than droplet transportation (<1 min).

The versatility of the Lo3DP platform is exemplified by its ability to automate temperature-dependent nucleic acid amplification assays, such as colorimetric LAMP. These assays offer an attractive alternative to qPCR by operating at a constant temperature and producing visible colorimetric outputs that can be interpreted without specialized instrumentation^26,27^. Despite their simplicity, LAMP workflows require precise liquid handling, controlled incubation at defined temperatures, and consistent optical readout—factors that traditionally limit scalability and reproducibility in resource-limited or decentralized settings. The Lo3DP addresses these challenges by integrating programmable reagent dispensing, on-chip thermal regulation, and image-based detection. This enables high-throughput screening and optimization of primer sets, accelerating assay development for a broad range of genetic targets.

While the current Lo3DP implementation supports key droplet operations such as transportation, merging, heating, and optical analysis, further expansion of its hardware and software ecosystem will broaden its scientific applicability. Integrating additional sensing modalities, such as fluorescence, impedance, or electrochemistry, would enable multiplexed and kinetic assays beyond colorimetry. From a software perspective, incorporating artificial intelligence (AI) and large language models (LLMs) could infuse adaptive intelligence into every step of lab automation: from microfluidic chip design and path planning to G-Code generation and real-time data interpretation. Acting as an “experimental co-pilot,” an LLM-enhanced Lo3DP could autonomously generate protocols, detect anomalies, and interpret outcomes, lowering the barrier for non-expert users. These advances, building on the Lo3DP architecture, can pave the way for accessible, democratized, and intelligent laboratory automation.

## V. Methods

### Materials for the Lo3DP System

The Lo3DP system was constructed using low-cost, widely accessible components, such as 3D-printer parts. The base motion system was derived from Ender-3 (Creality, Shenzhen, CN) and Prusa MK4S (Prusa Research, Prague, CZ) 3D printers. Custom connectors, holders, and other mechanical components were 3D-printed in-house using PLA filament (SUNLU, CN). The user interface was hosted on a Raspberry Pi 5 (8 GB), connected to a 10.1″ touchscreen monitor (ROADOM, CN) for GUI display and a 5″ touchscreen (ELECROW, CN) for the liquid-handling module. For optical imaging, a 64 MP Autofocus Camera (Arducam Technology, CN) was mounted on the manipulation head. Spectrophotometric measurements were performed using the AS7262 Visible Spectral Sensor (SparkFun Electronics, CO, USA). Magnetic droplet actuation used stacked Neodymium-Iron-Boron (NdFeB) magnets (1 mm thick each, >10 mm in total, 3 mm in diameter) and ferumoxytol (Ferraheme, AMAG Pharmaceuticals, MA, USA) as the magnetizer in the droplet. All microfluidic chips were filled with fluorinated oil (Novec 7500, 3M, MN, USA) containing 0.5 % biocompatible surfactant (Pico-Surf, Sphere Fluidics, NJ, USA). A standard piezoelectric voltage generator from a commercial fire lighter (BIC Multi-Purpose Party Lighter) provided the electrical pulses for contactless droplet merging.

### Microfluidic Chip Fabrication

Lo3DP microfluidic chips were fabricated by assembling multiple layers of transparent films (Apollo, LD Products, CA, USA), double-sided adhesive tapes (9474LE 300LSE, 3M, MN, USA), and an acrylic sheet (Acrylic United States, NY, USA). The multilayer stack consisted of a 1 mm acrylic sheet sandwiched between two 150 μm adhesive layers, with 100 μm transparent films on the top and bottom to enclose the microfluidic channels (**Fig. S5**). 2D microchannel patterns were designed in Autodesk AutoCAD and precisely cut on acrylic sheets using a laser cutter (Speedy 100, Trotec Laser, MI, USA). The transparent films were laser-cut to define the overall chip outline and inlet/outlet ports, then sequentially cleaned with 100% ethanol and deionized water. The cleaning process was repeated twice. When no surfactant was used, superhydrophobic coating was required to prevent droplet adhesion. The coating procedure involved: (1) treating the uncovered microfluidic chip (without the top transparent film) with NeverWet base coat and top coat (Rust-Oleum, USA), followed by 30 min of resting in the fume hood for superhydrophobic surface formation; (2) treating a separate transparent film with the same coating method; and (3) aligning and bonding the treated transparent film to the coated microfluidic chip to enclose the channels.

### Graphical User Interface (GUI)

An intuitive graphical user interface (GUI) was developed to enable user–Lo3DP communication and streamline microfluidic automation (**Fig. S1**). The GUI supports both the custom design of new workflows and the execution of existing biological assays. Communication between the GUI and the Lo3DP hardware occurs via a serial port, which sends G-Code that controls the manipulation head’s three-axis positioning with 0.1 mm resolution, enabling diverse functionalities. The interface also controls the print bed temperature, allowing programmable heating during assays. The GUI comprises four modular panels: (i) Editor (**Fig. S1A**) for manipulation-head navigation, (ii) Camera (**Fig. S1B**) for microfluidic chip imaging, (iii) G-Code Preview (**Fig. S1C**) for command generation and visualization, and (iv) Syringe Setup (**Fig. S1D**) for automated liquid handling. This structure provides coordinated control of droplet movement, reagent dispensing, heating, assay procedure timing, and image acquisition. The GUI was implemented in Python using the PySimpleGUI and Tkinter libraries. Additional details and workflow configurations are provided in **Note S4**.

### Transportation Speed Measurement

To characterize the maximum speed at which ferrodroplets could be reliably tracked, we conducted measurements using a microfluidic chip with a 50 mm × 10 mm × 1 mm chamber. The chip was positioned at the center of the Lo3DP print bed, and the stacked magnets were lowered until the bottom surface was ∼0.2–0.5 mm above the chip surface, without making physical contact. Ferrodroplets ranging from 0.1 μL to 100 μL in volume and containing 0.5% to 15% ferumoxytol were characterized (**Table S4**). For each test, a single ferrodroplet was magnetically guided along a 50-mm straight trajectory by displacing the stacked magnets via G-Code. The magnet velocity was controlled by adjusting the feed rate (F value) in the G-Code. A trial was considered successful if the droplet remained intact and followed the stacked magnets for >45 mm. Starting from a low transportation speed, like 0.5 cm/s, the speed was increased in 0.5 cm/s increments. Testing continued until the droplet either failed to follow the stacked magnets or broke apart. The highest speed at which the droplet completed a successful follow-up was recorded as its maximum trackable velocity. To compare ferrofluid droplets with particulate suspensions, we performed parallel measurements on droplets containing 50% magnetic beads (AMPure XP # A63880, Beckman Coulter, ∼1 μm diameter, 0.5–10 μL droplet volumes), following a similar procedure.

### Long-term Transportation Validation

To evaluate the long-term stability of ferrodroplet transportation on the Lo3DP platform, we fabricated a microfluidic chip with a 40 mm × 10 mm × 1 mm chamber. We loaded a 10 µL, 2.5% ferrodroplet into the oil-filled channel. The chip was positioned directly above a mini-spectrophotometer placed on the print bed, using a custom chip–sensor alignment fixture (**Fig. S6**) that centered the ferrodroplet over the spectrophotometer pinhole. An LED was placed adjacent to the chamber to provide constant illumination. During testing, the stacked magnets were programmed to drive the ferrodroplet back and forth across the top area of the pinhole, causing periodic occlusion and recovery of the light path. This motion was repeated continuously at 0.4 cycles per second for a total duration of 40,000 s, while the spectrophotometer recorded the resulting light-intensity fluctuations.

### Droplet Generation with Target Volumes

Target-volume droplets were generated using two microfluidic designs, a hook-based dispenser and a nozzle-based droplet generator, optimized for different volume ranges. For the hook-based dispenser, droplet volume was determined by the cross-sectional area of the hook structure. By incrementally increasing this area, we produced daughter droplets ranging from 0.5 to 9 μL. The larger hook structures were suboptimal due to inconsistent droplet splitting. This approach was validated for 2.5% and 5% ferrofluid formulations.

To generate larger droplets, a nozzle-based droplet generator was utilized (**Note S5**). Using this design, the stacked magnets were positioned directly above the nozzle neck to draw out a portion of the parent ferrodroplet. The daughter droplet was pinched off by moving the magnet laterally. For this structure, droplet volume was controlled by the pause time (**Δt**) when the magnets were held over the nozzle. Calibration experiments were conducted for 2.5% and 5% ferrodroplets to establish the quantitative relationship between **Δt** and droplet volume. Using these calibration curves, we reproducibly generated daughter droplets ranging from 10 to 25 μL by adjusting the appropriate **Δt**. Droplet volumes for both microfluidic designs were quantified from images acquired by a digital camera. The cross-sectional area of each droplet was measured in ImageJ and converted to volume using a pre-established calibration between droplet footprint and volume. Together, the hook-based and nozzle-based designs enabled a combined dynamic droplet generation range of 0.5–25 μL.

### Experiment and Indexing of Droplet Mixing

To evaluate the mixing performance on the Lo3DP, we merged 2 μL of dyed droplets with 8 μL of 2.5% ferrodroplets. Immediately after merging, the stacked magnets performed rotational stirring over the droplet. We systematically varied two parameters: the number of rotational cycles (0, 10, 20, or 30) and the rotation radius (0, 0.5, 2.5, or 3.5 mm) with a fixed speed of 0.5 cm/s. These combinations yielded 10 experimental conditions to assess how rotational motion influences mixing. After each run, droplets were imaged using a digital camera under uniform illumination. Images were circularly cropped to isolate the droplet and converted from RGB to HSV color space for better segmentation of the dyed region. The *Mixing Index* (MI) was computed based on the fraction of the dyed area within the total area. Specifically, the number of pixels falling within the predefined hue range corresponding to the dye was divided by the total number of droplet pixels (Python code for mixing efficiency analysis is available on GitHub at https://github.com/xxxx). An MI approaching 1.0 indicated near-complete color homogenization, whereas lower values reflected incomplete mixing. These indices were used to compare the relative efficacy of different rotational radii and cycle counts, with the static mixed control droplet serving as the reference.

### Magnetic Bead Separation

To evaluate the ability to separate magnetic beads (MBs) (AMPure XP # A63880, Beckman Coulter), we fabricated a microfluidic chip containing a 0.5-mm-radius semicircular chamber designed to trap MBs exclusively. 2.5% ferrodroplets were prepared with 10%, 50%, or 100% magnetic bead suspensions (∼1 μm diameter) and introduced into the chip. The stacked magnets guided each droplet along a predefined path and drew it against the semicircular chamber wall, where the magnetic field concentrated and immobilized the beads to form a cluster. After holding the droplet in place for several seconds, the stacked magnets were quickly moved away to extract the 2.5% ferrodroplet from the chamber, leaving the trapped MB cluster in place. To quantify separation efficiency, droplets were imaged before and after MB removal using a Nikon Ti-E inverted microscope to capture 16-bit light-transmission images that were then analyzed using ImageJ. Light-intensity values were compared with a 0% bead reference control droplet to determine the completion of MB removal; post-separation images with transparency approaching or equal to that of the reference 2.5% ferrodroplet indicated near-complete or complete separation of MBs.

### Automated Serial Dilutions with Lo3DP

Automated serial dilution experiments were conducted using three variations of a microfluidic design, each containing dispenser geometries matched to the target dilution factor. For 2× dilutions, the chip incorporated paired 5 μL buffer and 5 μL sample dispensers; for 5× dilutions, 8 μL buffer and 2 μL sample dispensers; and for 10× dilutions, 9 μL buffer and 1 μL sample dispensers. To optimize the visualization of colorimetric changes and ensure that the dynamic range covered the dilution series, the initial dye concentration was adjusted for each experiment: 1/64× stock for 2× dilutions, 1/10× stock for 5× dilutions, and undiluted stock for 10× dilutions. For each dilution protocol, the Lo3DP was programmed to dispense the required buffer droplet, aliquot the sample droplet using the corresponding dispenser, merge the droplets via contactless electrofusion, and mix the merged droplet using magnetic stirring before proceeding to the next dilution cycle. Each workflow produced four sequential dilution cycles, as shown in **Fig. 4 B-D**.

Following completion of the automated dilution sequence, droplets from all dilution steps, including the starting sample droplets, were imaged using a digital camera under consistent illumination. Colorimetric quantification was performed by measuring the 1−*R ratio*, where *R ratio* denotes the ratio of the droplet’s measured intensity to that of the reference control droplet. Log-transformed (1−*R ratio*) values were used to plot dilution curves for each protocol. Parallel 10× dilution experiments were conducted using 2.5% and 10% ferrofluid to evaluate the impact of ferrofluid concentration on optical properties.

### Automated Colorimetric LAMP-on-Chip with Lo3DP

Colorimetric loop-mediated isothermal amplification (LAMP) assays were carried out on the Lo3DP platform using the WarmStart® Colorimetric LAMP 2X Master Mix (New England Biolabs). LAMP primer sets were designed using the NEB LAMP Primer Design Tool (v1.4.1) and synthesized by Thermo Fisher Scientific. Before conducting on-chip amplification, we calibrated the system’s thermal behavior by measuring the chip’s temperature as we incrementally adjusted the 3D printer bed setpoint (5 °C increments from 25 °C to 110 °C). A strong linear relationship between the programmed bed temperature and the chip temperature was established, enabling precise temperature control. Temperature stability at 65 °C was assessed across 10 independent trials, and the coefficient of variation (CV) was calculated to validate consistent isothermal incubation.

For on-chip LAMP reactions, droplets containing LAMP master mix, five candidate primer sets (**Table S5**), positive DNA templates, and negative controls were preloaded into separate chambers on the microfluidic chip. On the Lo3DP platform, droplets were magnetically transported, merged, and assembled into five parallel LAMP reactions, each containing a positive reaction and a negative control. Following 30 min of isothermal incubation at 65 °C, endpoint colorimetric images were acquired using a digital camera. Amplified droplets exhibiting a pink-to-yellow transition were classified as successful reactions, while droplets remaining pinkish contained a non-amplifying or inefficient primer set candidate. To confirm amplification outcomes, droplets from each reaction were collected and subjected to conventional gel electrophoresis. Gel images were used to verify the presence or absence of expected LAMP amplicons and to validate the on-chip classification of primer sets as efficient or inefficient.

### Gel Electrophoresis Validation of LAMP Products

Verification of colorimetric LAMP reactions was performed using agarose gel electrophoresis. LAMP generates a mixture of stem–loop DNA structures and concatenated repeats of the target sequence, producing a characteristic ladder-like banding pattern on agarose gels. This method was used to validate BRCA1 amplification outcomes from on-chip reactions. End-point ferrodroplets containing LAMP products were analyzed on 1% agarose gels stained with GelRed nucleic acid dye. For each reaction, a 10 μL ferrodroplet was mixed with 6× TnTrack DNA loading dye and loaded into individual wells alongside a GeneRuler DNA ladder for size reference. Electrophoresis was performed at 130 V for 40 min, and gels were imaged using a Bio-Rad Universal Hood II imaging system. The presence of a distinct multi-band ladder pattern confirmed successful amplification, whereas its absence indicated inefficient or no detectable amplification.

## Supporting information

Supplementary Materials (Including Video Links)

## VI. Acknowledgments

This research is supported by the National Science Foundation Engineering Research Center (NSF ERC 1648451), the National Institutes of Health (NIH R01HL148182 and R01HL169695), and the U.S. Department of Veterans Affairs (I01-CX001901). We thank all collaborators (Jiarui Cui, Dr. Rajash Ghosh, Kiarash A. Sabet, Dr. Sam Emaminejad, Dr. Jeffrey J Hsu, Dr. Caiming Li, and Yuhang Li), mentees (Maximilian Mitterberger, Tianyi Li, Yuning Song, Chaidhat Chaimongkol, Sarah Taylor, Kyra Sunil, Damaris Cruz Alvarado, Tristan Lovely, Aiden BeGole, and Margherita Scussat), and visiting scholars (Yifei Tang and Binbin Zeng) for their dedication and contributions to this project.

## VII. Conflict of Interest

D.D., S.Y., Z.G., A.G., V.R. and CR.M. have a pending patent application on the Lo3DP technology. Other authors declare no conflict of interest.

## Notes

### Competing Interest Statement

D.D., S.Y., Z.G., A.G., V.R., and CR.M. have a pending patent application on the Lo3DP technology. Other authors declare no competing interest.

