## Supplementary Materials (Including Video Links) for "Lab-on-a-3D-Printer for Democratized Reconfigurable Digital Microfluidics"

**The PDF file includes:**

**Note S1.** Colorimetric and fluorescent properties of ferrodroplets
**Note S2.** Dispensing uniformity across ferrofluid concentrations
**Note S3.** Principle and step-by-step workflow of serial dilution on the Lo3DP
**Note S4.** Technical details of the graphical user interface (GUI)
**Note S5.** Variations of droplet generation nozzles across ferrofluid concentrations
**Table S1.** Applications of repurposed 3D printers for lab automation
**Table S2.** Comparison of robotic and microfluidic automated liquid handling systems
**Table S3.** Low-cost custom-built liquid handling systems
**Table S4.** Transportation speed across ferrofluid concentrations
**Table S5.** Sequence of LAMP primers
**Fig. S1**. Graphical user interface (GUI)
**Fig. S2** Step-by-step illustrations of the droplet generation and bead separation
**Fig. S3.** Colorimetric and fluorescent properties of ferrodroplets
**Fig. S4.** Original gel electrophoresis image of the colorimetric LAMP products
**Fig. S5.** Fast prototyping pipeline of microfluidic chips
**Fig. S6.** Spectrophotometric sensor fixture
**Fig. S7.** Dispensing uniformity across ferrofluid concentrations
**Legends** for movies S1 to S6
**References** (1 - 25)

**Other Supplementary Materials for this manuscript include the following:**

**Movie S1** (.mp4 format)**.** Droplet dispensing (0.5 - 5 μL)
**Movie S2** (.mp4 format)**.** Dispensing uniformity across ferrofluid concentrations
**Movie S3** (.mp4 format)**.** Nozzle-based droplet generation (10 - 25 μL)
**Movie S4** (.mp4 format)**.** Magnetic bead separation
**Movie S5** (.mp4 format). 2×, 5×, and 10× serial dilution
**Movie S6** (.mp4 format)**.** Automated LAMP assay preparation (Steps 1 and 2)

**Note S1.** Colorimetric and fluorescent properties of ferrodroplets

Colorimetric and fluorescent measurement experiments are typically performed with optically transparent fluids. Ferrofluids comprising iron oxide nanoparticles introduce optical scattering and absorption. To achieve droplet mobility while minimally compromising their optical properties, we investigated the effect of ferrofluid (FF) concentration (*i.e.*, 0.0, 0.5, 1.0, 2.5, 5.0, 7.5, and 10.0%) on signal decreases in colorimetric and fluorescent modalities. For colorimetric readouts, a 6-channel spectrophotometric sensor (AS7262 Visible Spectral Sensor, SparkFun Electronics, CO, USA) was used to evaluate how FF concentrations affect light transmission (negatively correlated with optical density) across several wavelengths of 450, 500, 550, 570, 600, and 650 nm. Fluorescence quantification was performed using a plate reader (BioTek Cytation 3, Agilent, CA, USA), and the readouts of fluorescein-laden droplets (1 µM, 0.1 µM, and 0.01 µM) at different FF concentrations were recorded. The detailed experimental procedures for the two modalities and the resulting signal decreases are as follows.

***S1-1. Dynamic range decrease for the colorimetric modality***

To investigate the decrease in dynamic range of the colorimetric readout of ferrodroplets across wavelengths of 450, 500, 550, 570, 600, and 650 nm, we developed a Python program (Python code for colorimetric property characterization is available on GitHub at https://github.com/xxxx) to acquire data from the sensor at 0.2-second intervals. The microfluidic chip was placed on top of the sensor and preloaded with oil and 25 µL droplets with FF concentrations of 0.5, 1.0, 2.5, 5.0, 7.5, and 10.0%. For each combination of wavelength and FF concentration (36 conditions in total), the height of the light source was adjusted until the signal readouts fluctuated around the maximum value from when the sensor read a Phosphate-buffered saline (PBS) droplet (0.0 % FF). 30 readouts were recorded and averaged as the maximum dynamic range at each channel. Then, the ferrodroplet was dragged to occlude the light path atop the sensor pinhole. 30 readouts were recorded, and the normalized dynamic range was calculated by dividing the average of them by the maximum dynamic range calculated above. **Fig. S3A** shows that normalized dynamic ranges for all channels decreased with increasing FF concentrations.

The 600 nm (orange) and 650 nm (red) channels were robust to increases in FF concentration and still maintained above 65% dynamic range. However, the readouts of the other four channels (450, 500, 550, and 570 nm) decreased dramatically with FF additions, especially at 450 nm (violet) and 500 nm (blue). When the FF concentration was 5.0%, the remaining dynamic range was below 25%. In contrast, with 0.5% FF droplets, over 68% of the dynamic range across all channels was retained, and with 2.5% FF droplets, all channels except 450 nm (violet) had dynamic ranges over 47%. Since our system uses stacked magnets that generate strong magnetic fields, ferrodroplets with 0.5%~2.5% FF concentrations can be easily moved (**Table S4**). These findings demonstrate a clear advantage of our system over the PCB-enabled ferrobotic system when working with low-FF droplets that have optimal optical properties for maximizing colorimetric readout signal.

#### *S1-2. Signal intensity decrease for the fluorescent modality*

We performed fluorescence characterization of droplets containing combinations of fluorescein (*i.e.*, 0.00 μM, 0.01 μM, 0.1 μM, and 1.0 μM) and ferrofluids (*i.e.*, 0.0, 0.5, 1.0, 2.5, 5.0, 7.5, and 10.0%) (28 conditions in total), as shown in **Fig. S3B**. Aliquots of 15 μL from each solution were pipetted into a 384-well plate, which was measured for fluorescence using a plate reader. The excitation wavelength was set to 485 nm, and the emission was recorded at 517 nm. Blank wells containing the corresponding FF concentrations, with 0.0 μM fluorescein, were used as the baseline shown as a grey solid line in **Fig. S3B**. The results revealed that at FF concentrations below 2.5%, fluorescence intensities were clearly distinguishable between 0.00 μM and 0.01 μM fluorescein concentrations. However, at concentrations above 5.0 % FF, the 0.01 μM dye solution showed signal blockage, making it indistinguishable from the background noise. The 5.0% FF concentration was a transition point, with higher FF concentrations resulting in substantial signal attenuation.

#### *S1-3. Optimal ferrofluid concentration: 2.5%*

Across both colorimetric and fluorescent characterization (**Fig. S3** and **Table S4)**, 2.5% ferrofluid provided the best balance between optical clarity and magnetic mobility. Lower concentrations provided excellent transparency but reduced magnetic responsiveness, whereas higher concentrations (>5.0%) caused substantial signal attenuation that obscured low-intensity colorimetric and fluorescent readouts. At 2.5%, ferrodroplets remained highly mobile under magnetic actuation while preserving sufficient optical transparency for quantitative measurements. This concentration was therefore selected as the standard condition for all Lo3DP assays.

### Note S2. Dispensing uniformity across ferrofluid concentrations

A hook-based microfluidic dispenser provides a repeatable means of dispensing ferrodroplets over a range of concentrations (**Movie S2**). The droplet splitting event is governed primarily by the volume of the hook structure and the movement path of the guiding magnet. To validate this concentration-independent performance, a total of 108 droplets were dispensed across six FF concentrations (*i.e.*, 2.5, 5.0, 7.5, 10.0, 12.5, and 15.0%), with 18 droplets imaged and analyzed under each condition. The dispensed droplets remained tightly centered around a target volume of approximately 1.00 μL, yielding an overall coefficient of variation of 3.88% (**Fig. S7**). Representative droplets across all concentrations exhibited identical sizes, shown in the inset images of **Fig. S7**, confirming that the uniformity of the dispensing mechanism was independent of magnetic responsiveness. This level of consistency supports its use in workflows that require precise droplet dispensing.

**Note S3.** Principle and step-by-step workflow of serial dilution on the Lo3DP

The Lo3DP performs serial dilutions by combining two precisely engineered dispenser structures—one for buffer droplets and one for sample droplets. Each dispenser produces a fixed droplet volume defined by its cross-sectional geometry. By selecting appropriate combinations of sample and buffer volumes (*e.g.*, 5 µL + 5 µL for 2×, 2 µL + 8 µL for 5×, or 1 µL + 9 µL for 10×), the system can generate a wide and tunable range of dilution factors. Because the dispensers can be fabricated across a continuum of droplet volumes (*e.g.*, 0.5–9.5 µL), the Lo3DP can theoretically produce dilution ratios ranging from 2× to 20× on a single microfluidic chip. Once the volumes are defined, the system executes a programmed sequence of dispensing, merging, and magnetic stirring to produce each dilution step in an automated, repeatable manner. Below, we provide detailed procedures for 2× (***S3-1***) and 5×/10× (***S3-2***) serial dilution (**Movie S5**), illustrating how these principles are implemented on our platform.

***S3-1. 2× serial dilution procedure***

1) Position the serial dilution microfluidic chip at the center of the print bed of the Lo3DP.
2) Start the program, load the experiment G-code, and click run.
3) The Lo3DP pipettes a 25 µL buffer droplet into the bottom-left buffer reservoir and a 10 µL sample droplet into the top-left chamber. Then, the system aligns the stacked magnets over the buffer reservoir.
4) The stacked magnets draw the 25 µL buffer droplet out of its well.
5) The buffer droplet is moved diagonally to the first dispenser, dragged into the dispenser, and then split. This results in a 5 µL buffer droplet nestled in the first dispenser.
6) Step 5 is repeated three times for the 2$^{nd}$, 3$^{rd}$, and 4$^{th}$ dispensers, resulting in three 5 µL buffer droplets nestled in the dispensers. The leftover 5 µL buffer droplet is moved into the top-right chamber for background control.
7) The four buffer droplets in the dispensers are sequentially dragged into mixing chambers on the top of the chip (*e.g.*, 1$^{st}$ dispenser buffer droplet to the 1$^{st}$ mixing chamber (numbered from the left) and 2$^{nd}$ dispenser buffer droplet to the 2$^{nd}$ mixing chamber).
8) The 10 µL sample droplet is dragged into the first chamber dispenser and split. The free 5 µL droplet is returned to the sample chamber, and the 5 µL droplet nestled in the chamber dispenser is moved to the 1$^{st}$ mixing chamber.
9) The droplets in the mixing chamber are merged via electrofusion, and the stacked magnets stir the unmixed droplet 30 cycles using a clockwise circular rotation.
10) Steps 8 & 9 are repeated three times in the 2$^{nd}$, 3$^{rd}$, and 4$^{th}$ mixing chambers, splitting the respective fully mixed droplet and moving the dispensed droplet into the next mixing chamber.
11) The final 10 µL droplet in the 4$^{th}$ mixing chamber is split following Step 8, but the 5 µL droplet nestled in the chamber dispenser is moved into the hook at the base of the right-most well instead of into the control droplet chamber.
12) The automated serial dilution is complete, and the stacked magnets move outside the chip.

***S3-2. 5×/10× serial dilution procedure***

1) Position the serial dilution microfluidic chip at the center of the print bed of the Lo3DP.
2) Start the program, load the experiment G-code, and click run.
3) The Lo3DP pipettes 24/27 µL buffer droplet into the left buffer reservoir and a 16/18 µL buffer droplet in the right buffer reservoir, located at the bottom-left corner of the chip (all volumes listed for 5×/10×, respectively). Then, the system pipettes a 10 µL sample droplet into the top-left chamber. The system aligns the stacked magnets over the right buffer reservoir.
4) The stacked magnets draw the right 16/18 µL buffer droplet out of its well.
5) The right 16/18 µL buffer droplet is moved diagonally to the first dispenser, dragged into the dispenser, and then split. This results in an 8/9 µL buffer droplet nestled in the first dispenser.
6) The leftover 8/9 µL buffer droplet is moved into the 4th mixing chamber. The droplet in the first dispenser is dragged to the 3rd mixing chamber.
8) The stacked magnets align over the left 24/27 µL buffer well and draw it out of its well.
9) Step 5 is repeated twice for the 1st and 2nd dispensers, resulting in another two 8/9 µL buffer droplets nestled in the dispensers. The leftover 8/9 µL buffer droplet is moved into the top-right chamber for background control.
10) The buffer droplet in the 1st dispenser is dragged to the 2nd mixing chamber, and the buffer droplet in the 2nd dispenser is dragged to the 1st mixing chamber.
11) The 10 µL sample droplet is dragged into the first chamber dispenser and split. The free 8/9 µL droplet is returned to the sample chamber, and the 2/1 µL droplet nestled in the chamber dispenser is moved to the 1st mixing chamber.
12) The droplets in the mixing chamber are merged via electrofusion, and the stacked magnets stir the unmixed droplet 30 cycles using a clockwise circular rotation.
10) Steps 11 & 12 are repeated three times in the 2nd, 3rd, and 4th mixing chambers, splitting the respective fully mixed droplet and moving the dispensed droplet into the next mixing chamber.
11) The final 10 µL droplet in the 4th mixing chamber is split following Step 11, but the 2/1 µL droplet nestled in the chamber dispenser is moved into the hook at the base of the right-most well instead of into the control droplet chamber.
12) The automated serial dilution is complete, and the stacked magnets move outside the chip.

**Note S4.** Graphical User Interface (GUI) technical details

The GUI contains four control panels: Editor, Camera, G-Code Preview, and Syringe Setup. The panels can be navigated through the menu bar at the top of the GUI. The Python code is available on GitHub at https://github.com/xxxx.

***S4-1. Editor panel***

The Editor panel (**Fig. S1A**) provides a graphical representation of the print bed of the Lo3DP, the shape outlines of microfluidic cartridges, and the biological assay procedures. A grid on the GUI represents the print bed and supports dimensions between 40×40 $mm^2$ and 200×200 $mm^2$. The shape outline of microfluidic cartridges can be superimposed on the grid using the Import DXF button, which uploads the outline of the cartridge configuration using the Drawing Interchange Format (DXF). The user can then lock the location of the cartridge outline on the grid and generate an assay protocol by specifying the sequential movement steps of the entire assay based on the assay requirements and cartridge outline. The assay protocol is encoded into the navigation path of the manipulation head, consisting of multiple locations created by left-clicking on the grid or by manually entering location coordinates. Subsequent location points are automatically linked with each other, creating a single movement path. Since the manipulation head has multiple functional elements (*e.g.*, stacked magnets, camera, and pipettor), the user can select the element to be used at each location. The user can add a skip location point to the path by right-clicking on the grid or using an Add Skip button; the manipulation head will be raised prior to moving to the target location. Increased distance between the manipulation head and the cartridge ensures that the stacked magnets will not change the position of non-targeted ferrodroplets within the microfluidic cartridge during skip operations.

The menu bar above the grid contains various buttons to manage assay protocols. It includes New (to clear the grid from any assay procedures and cartridges), Open (to add an existing assay protocol to the grid), Save as (to save a generated assay protocol to a user-defined folder), Send via Serial (to directly send the generated assay protocol to the Lo3DP in G-Code format), Autohome (to home axes of the Lo3DP), and Import DXF (to import microfluidic cartridge outlines to the grid in the DXF format). To increase the assay throughput and perform repetitive or different experiments simultaneously, the user can import multiple cartridges to the same working plate with the Import DXF button.

The head selection button underneath the grid contains Stacked Magnets, Camera, and Pipettor, corresponding to the three key functional elements mounted on the manipulation head. The user specifies the element used at each location by using the drop-down selection function.

The interactive checkbox menu located beneath the grid provides control over various grid functions. This includes Show Grid (to show/hide the grid outline on the print bed), Realtime Mode (to move the manipulation head in real-time with each left click on the grid), Entire View (to zoom out to display the entire 200x200 mm² print bed), Lock DXFs (to lock cartridge positions on the grid), Toggle Zoom (to enable zooming in or out with left/right clicks on the grid), and Toggle Capture (to enable adding capture points for imaging with the camera). The capture points added when the Toggle Capture checkbox is selected will be processed after the user clicks the Take Image button within the Camera panel (**Fig. S1B**). This functionality allows the user to re-read assay results once the assay and automatic imaging are completed, which helps correct assay readout-related errors (*e.g.*, when the initial capture points specified in the assay protocol were incorrect or the cartridge was misaligned during the experiment).

Instead of directly left or right-clicking on the grid, a new location can be added to the path manually by specifying the X, Y, and Z coordinates (Z Move or Z Skip) of the new location, along with the manipulation head movement speed, and clicking the Add Move or Add Skip button to add a regular or skip location, respectively. Compared with clicking the grid, manual location entry allows the user to specify coordinates with 0.01 mm resolution, thereby enabling more precise sample transfer and manipulation within the microfluidic cartridge. In addition to adding a new location, the user can add a time delay (in milliseconds, e.g., 1000 ms) between subsequent movements by clicking the Add Delay button and specifying a delay time for incubation or image capture. This manual point addition panel is located beneath the checkboxes menu.

The Code Table, located at the top right corner of the Editor panel, displays the current navigation path of the manipulation head. Each entry in the table includes the movement type (Move, Skip, or Pause), the X, Y, and Z coordinates (in mm) of the corresponding point within the path (indicating where the manipulation head has moved to at this step), the pause time (in ms), and the movement speed (in mm/min). The Code Table also allows users to select and manually modify a point (*e.g.*, adjust its coordinates, movement speed, or pause time) and delete individual points.

Finally, the Debug Terminal at the bottom right corner of the Editor panel displays the manipulation head movements based on user inputs. It also logs each action the user has performed on the grid and reports any errors or warnings during system operations.

***S4-2. Camera panel***

The Camera panel (**Fig. S1B**) allows the user to specify major camera parameters such as exposure time (in seconds), camera-to-sample distance (Z height, in mm), and resolution. The user can also specify the number of images to capture per location (N) and the interval between subsequent captures (in seconds) to enable both endpoint (N=1) and time-lapse (N>1) image capture modalities. The Auto Focus button automatically adjusts camera focus, allowing clear, focused images to be obtained. The Take Image button initiates image capture based on the user-specified parameters and the capture points defined on the grid in the Toggle Capture mode. The Auto Focus and Take Image buttons are located underneath the input fields with user-specified camera parameters. Captured images are displayed on the left side of the Camera panel, and the user can navigate between the images using the Previous and Next buttons located beneath the displayed images. All captured images are also saved in a user-defined folder. We note that the Z height value in the Camera panel and the Z move value in the Editor panel represent the same value; therefore, updating one will automatically update the other.

#### *S4-3. G-Code Preview panel*

The G-Code Preview panel (**Fig. S1C**) displays G-Code commands generated based on user inputs from the Editor panel. The subsequent G-Code functions and corresponding descriptions are displayed line-by-line within the preview window.

#### *S4-4. Syringe Setup panel*

The Syringe Setup panel (**Fig. S1D**) enables automated liquid handling, including introducing samples and buffers into microfluidic cartridges. The panel contains input fields to specify syringe pump parameters, such as the syringe type (*e.g.,* 0.5 mL, 1 mL, or 3 mL syringe), operation volume (in μL), and flow rate (in μL/min, up to 1000 μL/min). If the flow rate exceeds 400 μL/min, the program automatically selects the 1 mL syringe model. The user can switch the syringe operation mode between push and pull options to support fluid injection and withdrawal. The panel also contains a syringe operation management menu bar, which includes the Run button (to directly send the specified syringe parameters to the syringe pump via G-Code), the Save As button (to save the specified syringe parameters as a local G-Code file), and the Reset button (to reset syringe parameters). This bar is located underneath the input fields with user-specified syringe pump parameters. Syringe parameters and their descriptions are displayed in the preview terminal as G-Code and code comments. The terminal is located underneath the syringe operation management menu bar.

**Note S5.** Variations of droplet generation nozzles across ferrofluid concentrations

The Lo3DP employs two nozzle geometries, a 1-mm narrow nozzle and a 2-mm triangular wide nozzle (**Fig. S2**), to accommodate changes in magnetic responsiveness associated with varying FF concentrations. Droplets with higher FF infusion (*e.g.*, 5.0% FF) exhibit a strong magnetic response, readily deforming and flowing through a nozzle structure. In contrast, lower-FF droplets (*e.g.*, 2.5% FF) respond weakly to magnetic actuation, resulting in reduced mobility and greater difficulty in overcoming geometric constraints like a narrow nozzle. To compensate for this weakened magnetic responsiveness, wider nozzles are employed to reduce the hydrodynamic barrier to droplet extrusion. Thus, 1-mm nozzles are paired with 5% FF droplets, and 2-mm nozzles are paired with 2.5% FF droplets, ensuring reliable and repeatable droplet generation across ferrofluidic compositions.

In addition to nozzle width, two structural and operational adjustments enable robust droplet pinch-off for ferrodroplets with weaker magnetic responsiveness. First, triangular sharp edges are incorporated into the 2-mm nozzle design, providing a geometrically favorable break point that promotes clean droplet neck rupture. Second, as shown in **Fig. S2** and demonstrated in **Movie S3**, an additional "pull-up" step is introduced. After positioning the stacked magnets above the nozzle, the manipulation head is positioned slightly upward by 1 mm before the lateral pinch-off. This maneuver elongates the droplet neck to facilitate rupture. It simultaneously engages the stronger magnetic field at the edge of the stacked magnet column, thereby improving traction for low-FF droplets. Together, these structural and operational strategies enable efficient extrusion and pinching of both high-FF (5.0%) and low-FF (2.5%) droplets, thereby maintaining volume consistency within the 10–25 μL range.

**Table S1**. Applications of Repurposed 3D Printers for Lab Automation

| **(A)** | ***3D Printers for Lab Automation*** | | | | |
|---|---|---|---|---|---|
| Ref. No | 3D Printer Name | Application(s) and Area(s) | Utilized Component(s) | Major Modification(s) of the 3D Printer | Advantage(s) |
| [1] | Creality Ender-3 | Tissue staining (Histology, Pathology); Surface modification and dip coating (Chemistry, Materials Science) | Both the printer itself and the 3D printed parts | A jar holder for keeping the staining jars on the printer bed;<br>A headpiece for mounting the slide holder | Cheap (~$215), modular, easy to set up, very programmable, consistent, versatile, reversible, open-source |
| [2] | D-Bot Core-XY | High-throughput time-lapse imaging for bacteria (Microbiology) | Both the printer itself and the 3D printed parts | A Raspberry Pi camera for colorimetric and fluorescent imaging;<br>A G-code control system with Arduino and Raspberry Pi | Cheap (<$700), customizable, open-source, versatile, remote control, high positional accuracy |
| [3] | WANHAO Duplicator i3 | High-resolution mass spectrometry imaging (Bioimaging, Histology) | Both the printer itself and the 3D printed parts | A 3D-printed block holding the nebulizer connected to a gas line and a 6-port HPLC valve | Cost-effective (~$2600), reproducible, consistent, high analytical accuracy |
| [4] | Printrbot Simple | Nucleic acid extraction and amplification (Molecular Diagnostics) | The printer itself | A custom heat block from the extruder heater for extraction;<br>A disposable tip comb holding a set of magnetic rods controlled by the extruder motor | Cheap (~$399), rapid, easy to set up, reversible, portable, versatile, sensitive, sample-to-answer |
| [5] | Prusa Mendell RepRap | Wax printing for prototyping paper-based microfluidics (Medical Devices) | Both the printer itself and the 3D printed parts | Five 3D-printed parts for assembling the wax extruder;<br>A heater made up of copper pipes and a nichrome heating element | Cheap (<$690), easy to set up, rapid, flexible, open-source |
| [6] | Creality Ender-3 | Dip coating of PMMA deposited on silicon substrates (Materials Science) | Both the printer itself and the 3D printed parts | A custom 3D-printed gripper to hold samples for dip coating;<br>An interpreter to translate dip coating parameters into G-code | Cheap (~$300-400, ~10x cheaper than commercial), reversible, open-source, customizable, superior film quality, consistent |
| [7] | Printrbot Play & Printrbot Simple | Molecular diagnostic sample preparation (Molecular Biology) | Both the printer itself and the 3D printed parts | A tip-comb attachment with magnets to conduct particle-based nucleic acid extraction | Cheap (~$400-750), open-source, rapid prototyping and customization, scalable |
| [8] | Trinus 3D | Gene sequencing library preparation (Molecular Biology) | Both the printer itself and the 3D printed parts | A combined pipette control & tip dispenser;<br>An electromagnetic system for moving plates;<br>An application-specific deck frame | Cheap (~$1000), open-source, rapid prototyping, high throughput |
| [9] | Prusa i3 RepRap | 3D printing of a chemical reaction device; Synthesis of Ibuprofen (Chemical Synthesis) | Both the printer itself and the 3D printed parts | A holder for polytetrafluoroethylene (PTFE)-lined dispensing needles for liquid handling, attached to the extruder | Cheap (<$800), Integrated fabrication and reaction execution, open-source, scalable, reproducible |

| *Table S1A Continues* | | | | | |
|---|---|---|---|---|---|
| [10] | RepRap | 3D printing of diagnostic devices; Antibacterial drug testing (Biomedical Device, Drug Development) | Both the printer itself and the 3D printed parts | Added ink-jet technology to dispense gels and handle small amounts of liquid | Cheap (~$500), Integrated diagnostics and therapeutics, portable, open-source, customizable |
| [11] | RepRap | Droplet characterization, Chemical evolution analysis (Systems Chemistry, Chemical Engineering) | Both the printer itself and the 3D printed parts | A liquid-handling syringe system replacing the extruder;<br>An integrated reagent mixing stage;<br>An integrated camera for droplet behavior imaging and analysis | Cheap (~$800), rapid prototyping, integrated imaging system, high-throughput, consistent, customizable |

| **(B)** | ***3D Printers for Liquid Handling*** | | | | |
|---|---|---|---|---|---|
| Ref. No | Name and Type of the 3D Printer | Key Parameters and Application(s) | Utilized Component(s) | Major Modification(s) of the 3D Printer | Advantage(s) |
| [12] | Creality Ender-3 | 5 µL/min - 2 mL/min; (General use; microfluidic droplet generation) | Both the printer itself and the 3D printed parts | Three sets of threaded rods, M5 nuts, and 5 × 5 shaft coupler;<br>14 3D printed parts, like motor mounts | Cheap (~$193), easy to set up, open-source, large dynamic flow rate range, multi-channel |
| [13] | Creality Ender-3 Pro | 10 µL - 100 µL; (General use) | Both the printer itself and the 3D printed parts | A single-channel pipette added to a 3D printed mount;<br>A stepper motor for driving the pipette plunger | Cheap (~$325), accessible, customizable, open-source |
| [14] | Creality Ender-3 | 0.5 µL - 1250 µL; (Life science assays) | Both the printer itself and the 3D printed parts | A pipet assembly in replacement of the extruder; 3D printed parts, like the deck base; A heating module | Cheap (~$400), open-source, easy to set up, customizable |
| [15] | Creality Ender-3 & Ender-3V2 | 10 mL BD syringe; (High-viscosity biomaterial printing) | The printer itself | A "Enderstruder" extruder consisting of a readily available syringe, a 3D-printed core frame, a syringe carriage, and a stock motor | Cheap (<$260), open-source, customizable, accessible |
| [16] | Hellbot Magna I | 3 - 20 µL; (Dual reagent dispensing; lateral flow immunoassays) | Both the printer itself and the 3D printed parts | Two NEMA stepper motors connected to a 3D Printer controller; 3D-printed dispensing tip (DT) holders; Two syringes coupled to DTs with 3-way valves | Cheap (~$600), open-source, flexible, customizable, efficient dual dispensing |

**Table S2.** Comparison of robotic and microfluidic automated liquid handling systems

| **Ref. No.** | **Technology** | **Footprint** | **Instrument Cost** | **Consumables Cost** | **Volume Range** |
|---|---|---|---|---|---|
| [14] | Robotic Liquid Handler (*e.g.*, Opentrons) | 87cm × 69cm × 84cm | $15,000 | ~$100-200 (estimated) | 1 μL - 1000 μL |
| [17] | Electrowetting-on-dielectric (EWOD) | 7.5 × 3 $cm^2$ | >$1,000 (estimated) | $100-500 (estimated) | 3 μL - 5 μL |
| [18] | Electret-induced polarization on droplet (EPD) | < 20cm × 20cm (estimated) | ~$100 for control system | <$1 | 500 nL - 1 mL |
| [19] | Mechanical actuation on surface (MAOS) | 34.5 cm × 48.0cm × 33.0 cm | >$2,000 (estimated) | ~$30 | 500 nL - 200 μL |
| Present work | Lab-on-a-3D-Printer (Lo3DP) | 55cm × 55cm × 50cm | <$500 (w/o accessories)<br><$800 (w accessories) | ~$10 | 0.1 μL - 100 μL |

**Table S3.** Low-cost custom-built liquid handling systems

| Ref. No | Name of the system | Key Parameters and Application(s) | Utilized Component(s) | Advantage(s) | Disadvantage(s) |
|---|---|---|---|---|---|
| [20] | OpenWork station | 1 µL - 1000 µL; (Hydrogel processing; 3D cell culture) | Modular frame system; motion controller; onboard computer; liquid handling unit, light module | Cheap (~$900), modular, open-source, customizable, consistent, reproducible | Basic coding needed, no onboard sensing, bulky setup, manual protocol configuration, high assembly complexity |
| [21] | FINDUS | 20 - 1000 µL; (Life science assays, enzyme kinetics, solid-phase peptide synthesis, automated dilution series) | Modular 3D-printed structure; low-cost microcontroller; stepper motor actuation; pipette-based dispensing; scriptable software interface | Cheap (<$400), open-source, easy to set up, customizable, consistent, programmable, high pipetting precision | Manual pipette unit exchange, low throughput, no built-in UI, slow |
| [22] | OTTO | 1 - 1000 µL; (qPCR sample preparation, general life science assays, automated serial dilutions) | Modular motion platform; open-source electronics; micropipette-based liquid handling; programmable control system | Cheap (~$1500), open-source, modular, easy to set up, accurate, consistent, customizable, unattended automation | Low throughput, not plug-and-play, calibration needed, tip-dependent errors |
| [23] | EvoBot | 100 µL - 20 mL; (Modular liquid handling; microbial fuel cell nurturing; chemotactic droplet experiments; OCT scanning; 3D printing of bioelectrodes) | Modular hardware platform; open-source electronics; syringe-based liquid handling; programmable control system | Cheap (<$500), modular, open-source, customizable, easy to set up, versatile, reconfigurable | Basic coding needed, limited user support, occasional recalibration, not plug-and-play |
| [24] | OpenLH | 2 - 720 µL; (Serial dilution, color mixing, pH tests, density experiments, yeast growth assays) | Commercially available robotic arm; programmable motion controller; visual block-based programming interface; syringe-based liquid handling unit | Cheap (~$380), easy to set up, open-source, compatible with standard labware, customizable | Limited precision, low throughput |
| [25] | Liquid-handling Lego robots | 0.25 - 720 µL; (STEM education, liquid handling demos, life science teaching labs, automated mixing and dilution, hands-on robotics for biology) | Modular Lego-based structure; onboard controller; motor-driven syringe unit; basic sensor module; programmable user interface | Cheap (~$385), modular, open-source, easy to set up, programmable, educational, compatible with standard labware | Limited precision, low throughput, calibration needed, basic sensing only |

**Table S4.** Transportation speed across ferrofluid (FF) concentrations

| | **Droplet Movement Max Speed (cm/s) Characterization** | | | | | | | | | | |
|---|---|---|---|---|---|---|---|---|---|---|---|
| Droplet Volume (µL) | 0.1 | 0.5 | 1.0 | 2.5 | 5.0 | 7.5 | 10.0 | 15.0 | 20.0 | 50.0 | 100.0 |
| Magnet Diameter (mm) | 3.0 | | | | | | | 6.0 | | 8.0 | |
| Max Speed (0.5 % FF) | 0.1 | 0.1 | 0.5 | 0.5 | 0.5 | 0.5 | 0.5 | 0.5 | 0.5 | 0.5 | 0.1 |
| Max Speed (1.0 % FF) | 0.5 | 1.0 | 1.0 | 1.5 | 1.5 | 1.0 | 1.0 | 1.5 | 1.5 | 1.5 | 0.5 |
| Max Speed (2.5 % FF) | 3.0 | 3.5 | 4.0 | 4.5 | 4.0 | 1.5 | 2.0 | 4.5 | 3.5 | 4.0 | 2.0 |
| Max Speed (5 % FF) | 6.0 | 7.0 | 6.0 | 11.0 | 10.5 | 8.5 | 8.0 | 7.0 | 6.5 | 5.5 | 5.0 |
| Max Speed (7.5 % FF) | 13.0 | 14.0 | 14.0 | 18.0 | 18.0 | 14.0 | 13.0 | 9.0 | 8.0 | 5.5 | 4.5 |
| Max Speed (10 % FF) | 16.5 | 18.0 | 18.0 | 18.0 | 18.0 | 18.0 | 17.0 | 11.0 | 11.0 | 5.5 | 4.0 |
| Max Speed (12.5 % FF) | 18.0 | 18.0 | 18.0 | 18.0 | 18.0 | 18.0 | 18.0 | 13.0 | 13.5 | 6.0 | 4.0 |
| Max Speed (15 % FF) | 18.0 | 18.0 | 18.0 | 18.0 | 18.0 | 18.0 | 18.0 | 18.0 | 18.0 | 6.0 | 3.0 |

The transportation speed characterization was conducted in an immiscible oil (Novec 7500 w/0.5% Picosurf).
The heights of the small (Ø 3.0), middle (Ø 6.0), and large (Ø 8.0) magnet stacks are 14, 13.5, and 13.2 mm, respectively.
18 cm/s is the upper limit speed for the testing gantry.

**Table S5.** Sequence of LAMP primers

| **Name** | **Sequence** |
|---|---|
| BRCA_F3_1 | GCCCAAGTGATGCTCTGG |
| BRCA_B3_1 | ACATAGTGTCCCCCTCAAGG |
| BRCA_FIP_1 | TACGCCTCTCAGGTTCCGCCGGCGTGGGAGAGTGGATT |
| BRCA_BIP_1 | GGCAGTTTGTAGGTCGCGAGGATCACGAGGATTCCCCCA |
| BCRA_LF_1 | AATACCCATCTGTCAGCTTCGG |
| BRCA_LB_1 | GAAGCGCTGAGGATCAGGAAG |
| BRCA_F3_2 | GCAGGCACTTTATGGCAAAC |
| BRCA_B3_2 | GAAACCCCACAGCCTGTC |
| BRCA_FIP_2 | TCTCTCGGGGCTCTGGATTGGTCCTCTTCCGTCTCTTTCCT |
| BRCA_BIP_2 | TGGCTCTTTCTGTCCCTCCCATCCAAGGGGCTACCGCTAA |
| BCRA_LF_2 | TGCCCCCGGATGACGTAA |
| BRCA_LB_2 | CCTTGATTTCGTATTCTGAGAGGCT |
| BRCA_F3_3 | GCAGGCACTTTATGGCAAAC |
| BRCA_B3_3 | AATTCCCGCGCTTTTCCG |
| BRCA_FIP_3 | TCTCTCGGGGCTCTGGATTGGTCCTCTTCCGTCTCTTTCCT |
| BRCA_BIP_3 | TGGCTCTTTCTGTCCCTCCCATCCAAGGGGCTACCGCTAA |
| BCRA_LF_3 | TGCCCCCGGATGACGTAA |
| BRCA_LB_3 | CCTTGATTTCGTATTCTGAGAGGCT |
| BRCA_F3_4 | TGGGTGGCCAATCCAGAG |
| BRCA_B3_4 | GAAACCCCACAGCCTGTC |
| BRCA_FIP_4 | CGCTAAGCAGCAGCCTCTCAGAGACGCTTGGCTCTTTCTG |
| BRCA_BIP_4 | TGGCTCTTTCTGTCCCTCCCATCCAAGGGGCTACCGCTAA |
| BCRA_LF_4 | ACAATCAGAGGATGGGAGGGA |
| BRCA_LB_4 | AGATAAATTAAAACTGCGACTGCGC |
| BRCA_F3_5 | TGGGTGGCCAATCCAGAG |
| BRCA_B3_5 | GAAACCCCACAGCCTGTC |
| BRCA_FIP_5 | CGCTAAGCAGCAGCCTCTCAGAGACGCTTGGCTCTTTCTG |
| BRCA_BIP_5 | TGGTTTCCGTGGCAACGGAAAAGAAGTCTCAGCGAGCTCAC |
| BCRA_LF_5 | ACAATCAGAGGATGGGAGGGA |
| BRCA_LB_5 | AGATAAATTAAAACTGCGACTGCGC |

### Supplementary Figures

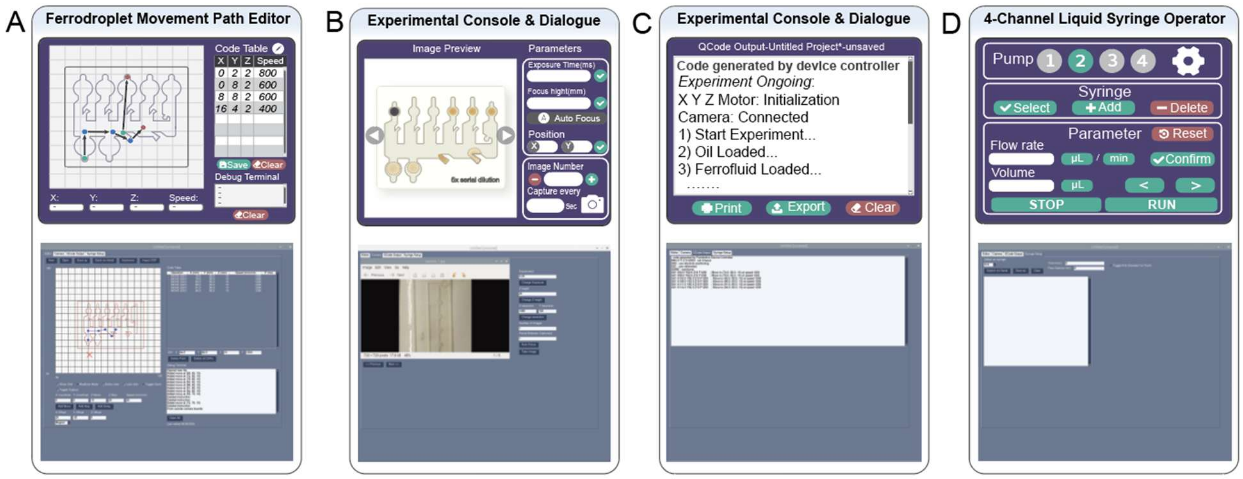


**Fig. S1.** Graphical user interface (GUI)

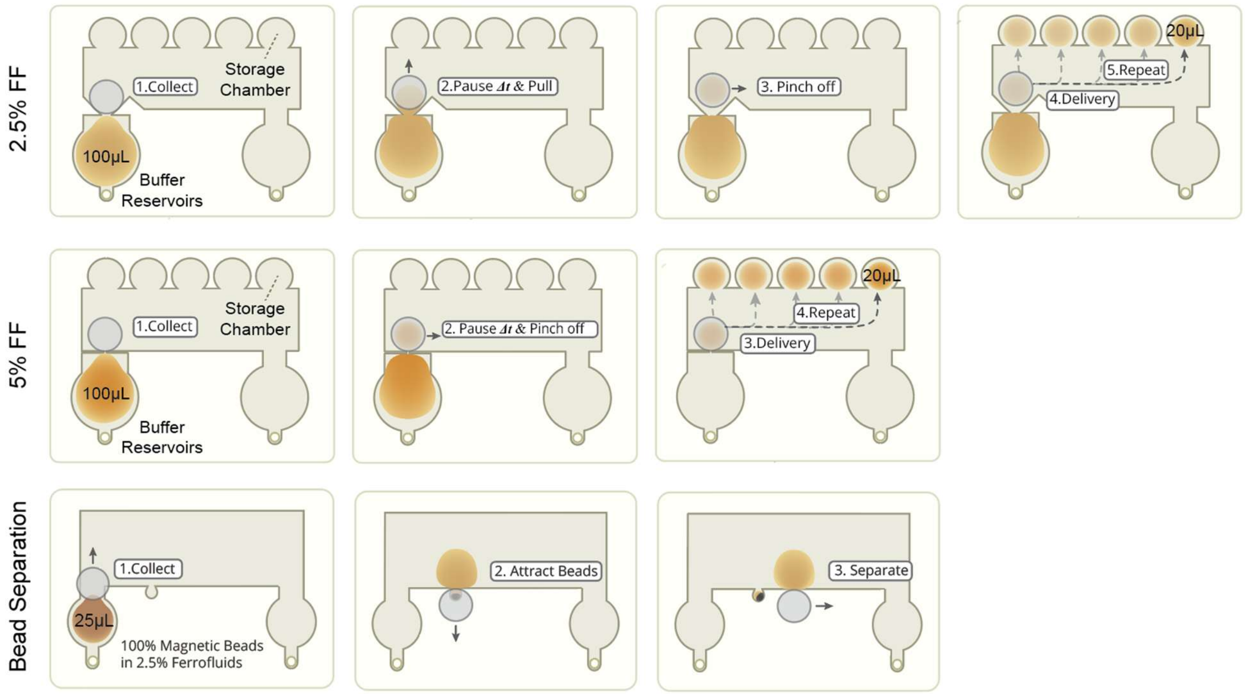


**Fig. S2.** Step-by-step illustrations of the droplet generation and bead separation

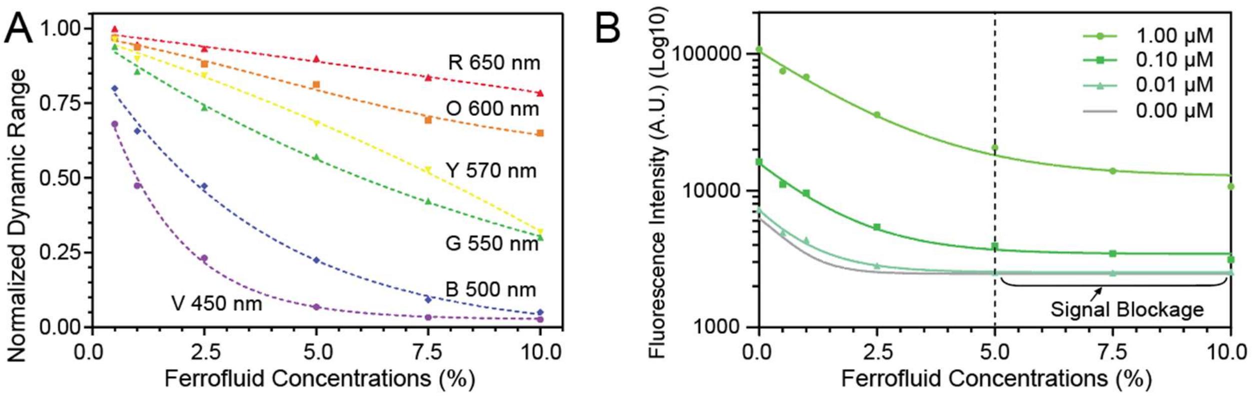


**Fig. S3.** Colorimetric and fluorescent properties of ferrodroplets

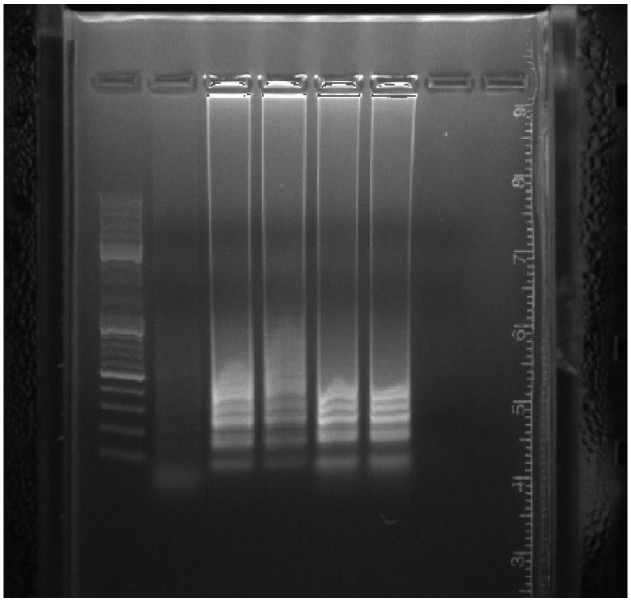

**Fig. S4.** Original gel electrophoresis image of the colorimetric LAMP products
(From left to right: 50-1500 bp Ladder, amplificons of LAMP primer sets 1 to 5)

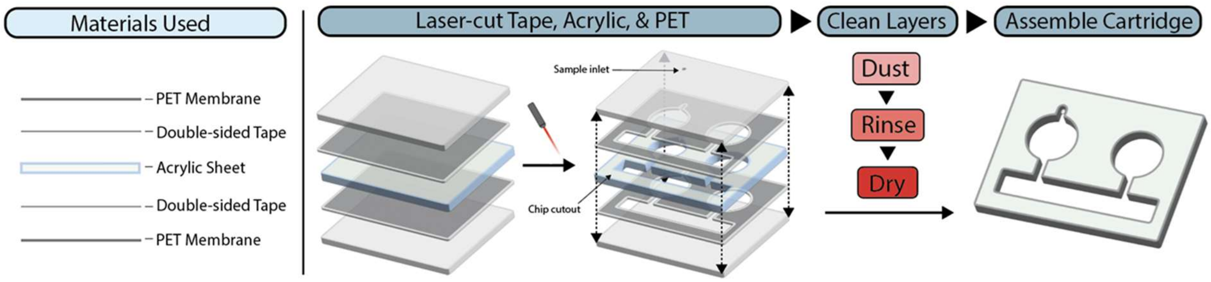


**Fig. S5.** Fast prototyping pipeline of microfluidic chips

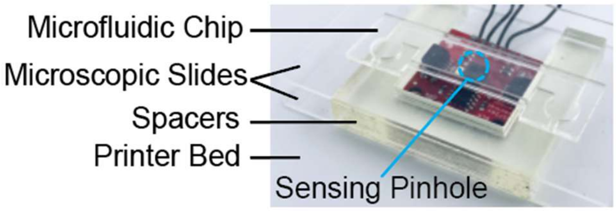


**Fig. S6.** Spectrophotometric sensor fixture

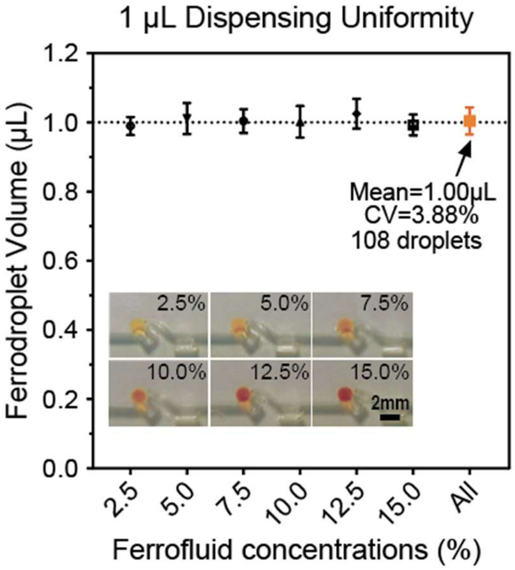


**Fig. S7.** Dispensing uniformity across ferrofluid concentrations

**Legends for movies S1 to S6**

**Movie S1** (.mp4 format)**.** Droplet dispensing (0.5 - 5 µL)
**Movie S2** (.mp4 format)**.** Dispensing uniformity across ferrofluid concentrations
**Movie S3** (.mp4 format)**.** Nozzle-based droplet generation (10 - 25 µL)
**Movie S4** (.mp4 format)**.** Magnetic bead separation
**Movie S5** (.mp4 format)**.** 2×, 5×, and 10× serial dilution
**Movie S6** (.mp4 format)**.** Automated LAMP assay preparation (Steps 1 and 2)

All the videos are available:
https://drive.google.com/drive/folders/1e2pgGo419E1fkUp2S3WThunx9F43ygVC?usp=sharing
